# Closed State Structure of the Pore Revealed by Uncoupled Shaker K^+^ Channel

**DOI:** 10.1101/2025.03.17.643777

**Authors:** Yichen Liu, Carlos Bassetto, Gustavo Contreras, Eduardo Perozo, Francisco Bezanilla

## Abstract

Voltage gated potassium (Kv) channels play key roles in numerous physiological processes from cellular excitability to immune response and are among the most important pharmaceutical targets. Despite recent advances in the structural determination of Kv channels, the closed state structure of strictly coupled Kv1 family remains elusive. Here, we captured the closed state structure of the pore in the Shaker potassium by uncoupling the voltage sensor domains from the pore domains. Ionic current, gating current and fluorescence measurements show that a conserved isoleucine residue in the S4-S5 linker region, plays a key role controlling the strength of the electromechanical coupling and the channel activation-deactivation equilibrium. Structural determination of completely uncoupled I384R mutant by single particle cryoEM revealed a fully closed pore in the presence of fully activated but non-relaxed voltage sensors. The putative conformational transitions from a fully open pore domain indicates a “roll and turn” movement along the whole length of the pore-forming S6 helices in sharp contrast to canonical gating models based on limited movements of S6. The rotational and translational movement posits two hydrophobic residues, one at inner cavity and the other at the bundle crossing region, directly at the permeation pathway, limiting the pore radius to less than 0.7Å. Voltage clamp fluorimetry of wild type channel incorporating a fluorescent unnatural amino acid strongly supports the cryoEM structural model. Surprisingly, the selectivity filter was captured in a noncanonical state, unlike the previously described dilated or pinched filter conformations. With the present experiment results, we propose a new gating model for strictly coupled Kv1 channels and the molecular mechanism of interactions among different functional states.

## Introduction

Voltage-gated potassium (Kv) channels regulate the passage of K^+^ ions across the cell by transducing the energy of transmembrane electric field. As the largest group of ion channels in the human proteome, Kv channels play key roles in cellular excitability, volume regulation, cell proliferation, immune response, and are among the most important pharmaceutical targets (Jan & Jan, 2012). Numerous structural studies on a variety of Kv channels have provided a great deal of information on the conformational landscape of the channels as a whole. However, given that the vast majority of the structures have been obtained in the nominal absence of an electric field, structural details of the closed state in “strictly coupled” Kv channels remain an unsolved problem in the field of ion channels.

In strictly coupled Kv channels, the electromechanical coupling (EMC) between the voltage sensing domain (VSD) and the pore domain (PD) is obligatory: the opening or closure of the channel requires activation or deactivation of the voltage sensors, respectively (Hirschberg et al., 1995; Islas & Sigworth, 1999; Sigg & Bezanilla, 1997). This is in contrast to a typical allosteric coupling, as in BK (Latorre et al., 2010), HCN (Flynn & Zagotta, 2018) or hERG (Bassetto et al., 2023; Lörinczi et al., 2015) channels for instance. While this tight coupling ensures a remarkably low “leak” potassium conductance at rest, it has also hindered our structural understanding of the gating mechanisms of strictly coupled Kv channels since at 0mV, the voltage sensors are typically observed in the activated (Up) conformation with the pore domain displaying an open inner bundle gate. As a result, the current gating model of strictly gated Kv channels, such as the Kv1 and Kv2 families, relies primarily on bacteria potassium channel structures (Doyle et al., 1998) and electrophysiological data (Holmgren et al., 1998; Tombola et al., 2006).

Here, we report the closed structure of the pore in a strictly coupled Kv1 channel, Shaker K^+^ channel, by disrupting the electromechanical coupling between the pore and the voltage sensors. A conserved isoleucine I384 in the S4-S5 linker region was identified as pivotal for the electromechanical coupling in Kv1 family (Haddad & Blunck, 2011). Single channel, ionic current, gating current and fluorescence measurements demonstrated that mutations to the side chain of I384 directly affects the strength of electromechanical coupling. In the most extreme case of I384R, the VSDs becomes completely uncoupled from the PD: the pore remained closed within a large range of voltages (-120mV to 180mV) while the VSDs activate/deactivate independently. Single-particle cryo-EM structure of the uncoupled channel revealed a collapsed permeation pathway, consistent with the expectation of the fully closed conformation. Site-directed fluorimetry measurements utilizing a fluorescent unnatural amino acid (UAA) strongly supports our structural observations. Additionally, structural rearrangements were also observed in the selectivity filter (SF) and the voltage sensors that likely underlie the structural basis for activation-inactivation coupling and hysteresis of the VSDs respectively. Taken together, we propose a new gating model for strictly coupled Kv channels and a molecular mechanism of interactions among different conformational states.

## Results

### Electromechanical coupling controlling residue in S4-S5 linker

A systematic mutagenesis survey of the S4-S5 linker, a region that has been demonstrated to be important for EMC (Isacoff et al., 1991; McCormack et al., 1991; Slesinger et al., 1993), led to the identification of a single residue that is able to tune the coupling strength between VSDs and PD. The conserved isoleucine 384 (Haddad & Blunck, 2011) locates at the N-terminus end of the S4-S5 linker and forms elaborate interactions with the intracellular end of the pore-forming S6 helix (Tan et al., 2022) (Fig. 1A, B). Replacing I384 with smaller residues such as alanine or cysteine, strengthened the already efficient electromechanical coupling (Fig. 1C and Supp. Fig. 1). I384C, for instance, activated in more negative voltages with much steeper voltage dependency, as illustrated by its conductance-voltage (GV) curve. More importantly, channel activation followed more closely the voltage sensor movement, measured in the gating-charge-voltage (QV) curve (Fig. 1 D). While in the wild type (WT) channels, the difference between the V_1/2_ for QV (measured with the nonconductive W434F mutant, Perozo et al., 1993) and GV curve was > 25mV (Fig. 1 E, F, Supp. Fig. 1, and Supp. Tables 1-2), that difference was less than 7.5mV in I384C, a hallmark for strengthened electromechanical coupling. On the other hand, mutating I384 to residues like glutamate, asparagine or leucine led to severe uncoupling of VSD from the PD (Supp. Fig. 1). This is characterized, in the case of I384L (Fig. 1 G), by large GV curve shifts (> 70mV) to more depolarized potentials, when compared to WT Shaker, together with a shallower slope in the GV curve (Fig. 1 H). Strikingly, while the GV was shifted to the right, the QV curve of I384L was shifted in the opposite direction (Fig. 1 H). Indeed, similar changes in the current kinetics and GV curves were also seen in hKv1.2 and hKv1.3 with mutations at equivalent position (I316 position in hKv1.2 and I386 position in hKv1.3), suggesting that I384’s critical role in the electromechanical coupling is conserved (Sup Fig. 3).

**Figure 1:**
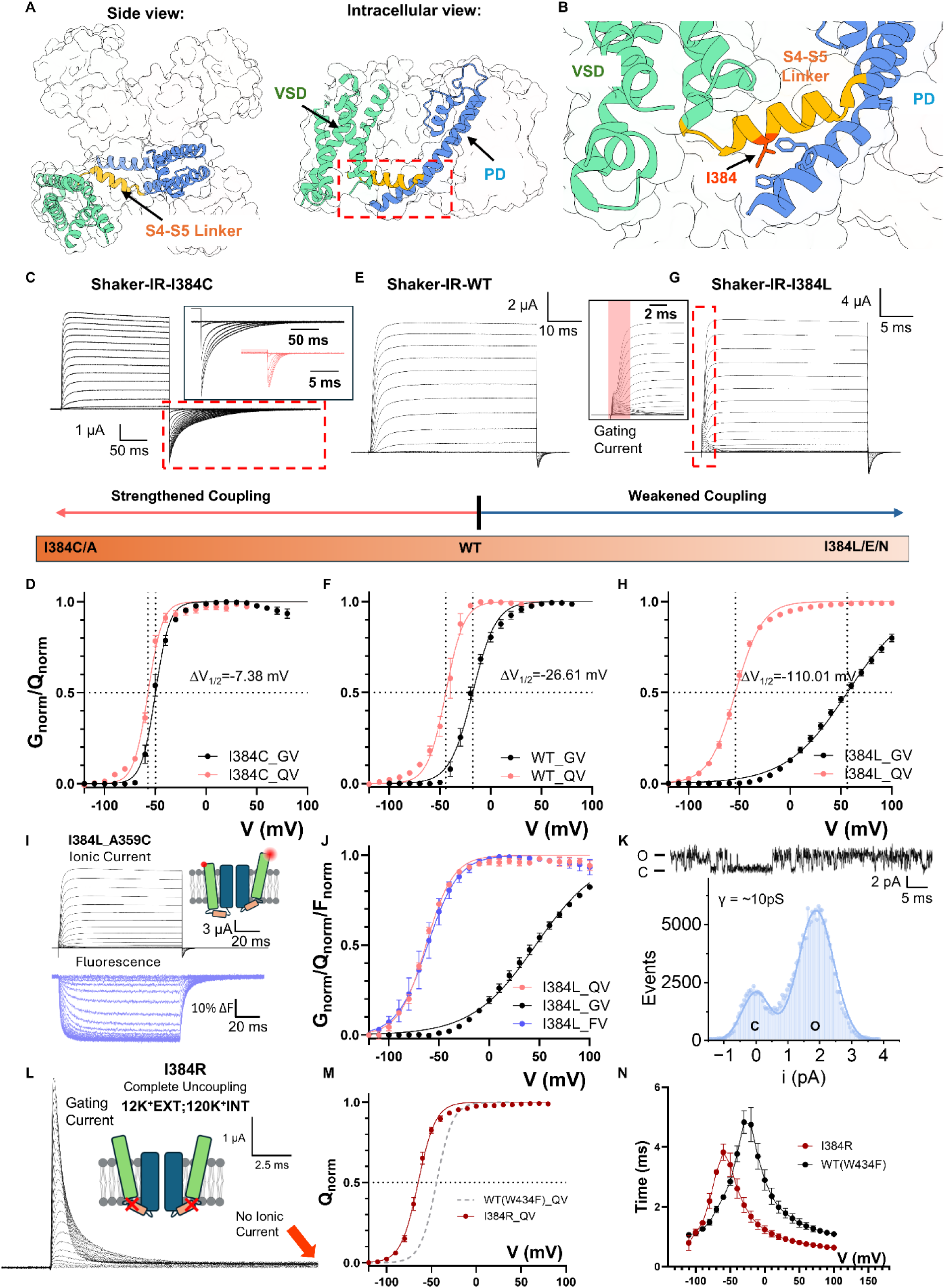
I384 residue controls the electromechanical coupling in Shaker potassium channel. **A)** Overall structure of Shaker potassium channel (PDB:7SIP). Shaker potassium channel is a homotetramer assembled in a domain-swapped manner. Each subunit has two major function domains, the <u>v</u>oltage <u>s</u>ensing <u>d</u>omain (VSD) and <u>p</u>ore-forming <u>d</u>omain (PD). Two function domains are connected by a short helix, the S4-S5 linker. **B)** Location of coupling controlling residue I384. I384 is surrounded by residues located in the S6 of the contiguous subunit. **C)** Representative ionic current traces from I384C. Inset in shows the comparison of the tail current between I384C in black and WT in red hyperpolarizing to the same voltage. Note the difference in time scale. Clearly the deactivation is much slower in I384C. **D)** GV and QV curves for I384C. The GV is much sharper and follows much closer the QV curve. **E)** Representative traces of WT Shaker channel. **F)** GV and QV curves for WT. **G)** Representative traces of I384L. The inset highlights the gating current resolved at the beginning of depolarizing pulse. GV and QV curves for I384L. The GV is significantly displaced to the right and is shallower. For **C)**, **E)**, **G)**, prepulse and returning pulse are both -120 mV and ΔV = 10mV. For **D)**, **F)**, **H)**, the data were fitted with a two-state model (details in method section) for easy visualization and calculation of change in ΔV_1/2_. Fitting results could be found in supplementary table 1 and 2. **I)** Ionic current and fluorescence traces from TMR-labeled at position A359C in I384L mutant background. Note there is no slow component in the fluorescence traces. **J)** Comparison of GV, QV and FV curves of I384L. The FV curve is essentially overlapping with the QV curve. **K)** One example of single channel currents. The single channel conductance is not significantly altered but flickering behaviors increased. **L)** Representative traces of I384R. Even in the presence of high K^+^ concentration in and outside of the cell no ionic current is detected, and only gating current could be seen. **M)** QV curve for I384R. Compared to the WT, the QV is shifted more than 30mV to the left. **N**) Time constants of on-gating kinetics of I384R and WT with W434F background. Clearly the gating current in I384 is faster. All the data is shown as Mean ± SEM. N = 3-6 independent experiments.

Since V_1/2_ gap between GV and QV is so large in the I384L, gating currents were easily resolved in the presence of ionic currents (Fig.1 G inset). To confirm that no additional slow component in the VSD movements was present in I384L that is responsible for the opening of the pore, tetramethyl rhodamine (TMR) was introduced at A359C in the extracellular loops of the VSD so that the movement of the voltage sensors would be evaluated via its fluorescence signal (Cha & Bezanilla, 1997; Kalstrup & Blunck, 2013; Mannuzzu et al., 1996). The fluorescence records showed no additional component in the VSD movement, and the voltage-dependent fluorescence (FV) curve fully overlaps on the QV curve (Fig. 1I, J). This result demonstrates that our gating current measurement reflect the true movement of VSDs. To confirm the observed effects were indeed a consequence of impaired coupling rather than due to changes in single channel properties, we performed noise analysis and single channel recordings (Alvarez et al., 2002; Sigworth, 1980) in I384L mutant. Noise analysis demonstrated that at 195mV, the maximum opening probability was less than 70% (Supp. Fig. 3) and single channel recordings showed that the unitary conductance level in I384L (Fig. 1K, ∼10pS, Supplementary Tables 3-4) was similar to the WT (∼12pS) (Alvarez et al., 2002), but flickering behaviors were increased (Fig. 1K and Supp. Fig. 3).

While in I384L, the impairment in the coupling was severe, we discovered another mutant that completely uncouples the voltage sensors from the pore, I384R. Despite the presence of K^+^ ions, I384R showed no ionic conduction, and only gating current could be recorded (Fig. 1L). The gating current itself activated at more hyperpolarized voltages compared to WT (Fig. 1M) and was kinetically faster compared to the WT with W434F background (Fig. 1N), as if an energetic load was taken off from the voltage sensors. This is also consistent with left-shifted QV curves seen in other uncoupled mutants such as I384N, I384L and I384E (Supp. Fig. 1, and Supp. Tables 1-2). In I384R, our recordings showed no discernible K^+^ current from -120mV to +180mV, demonstrating an exceptional structural stability of the closed pore. Additionally, the left-shifted QV curve allows the VSDs to transition more easily to the up conformation. As a result, at 0mV, I384R possesses a stably closed pore and four activated VSDs with little structural heterogeneity, an appealing target for structural investigation.

### Closed state structure revealed by a completely uncoupled channel

We expressed, purified and solved the structure of Shaker-IR-I384R, by single-particle cryo-EM. The structure was globally resolved to 3.5 Å, with clear densities for all the transmembrane helices (Fig. 2A, B, Supp. Fig. 4 and 5). Similar to previously determined structures, the uncoupled channel assembled as a domain-swapped homotetramer (Fig. 2A, B) (Tan et al., 2022). However, unlike the WT channel structure captured with an open pore, the uncoupled channel clearly displayed a collapsed permeation pathway. (Fig. 2C, D). Compared to the open conformation of Shaker, the pore-forming S6 helices in the closed pore underwent a “roll and turn” movement. The translational movement towards the permeation pathway brought the backbone of the S6 helices closer (Fig. 2C), and a rotational movement posited the hydrophobic I470 in the inner cavity and V474 near the bundle crossing directly into the pore where the two narrowest points of the closed pore are located (Fig. 2D). Pore radius utilizing HOLE calculation showed a closed permeation pathway with radius less than 0.7 Å (Smart et al., 1996), leaving ion conduction an impossibility (Fig. 2E, F). The rotating-in of the I470 residue led to a total collapse of the water-filled inner cavity underneath the selectivity filter, an unexpected result (Fig. 2E, F). Current gating model predicts a “hinge-like” movement for the channel activation where a kink is created in the middle of the S6 helices around the conserved PVP motif and the intracellular half of the helices crosses or separates to close or open the channel (Del Camino et al., 2000; Del Camino & Yellen, 2001; Holmgren et al., 1998; Liu et al., 1997; Tombola et al., 2006; Webster et al., 2004). Thus, the observed movement around the upper part of the S6 helices was surprising.

**Figure 2:**
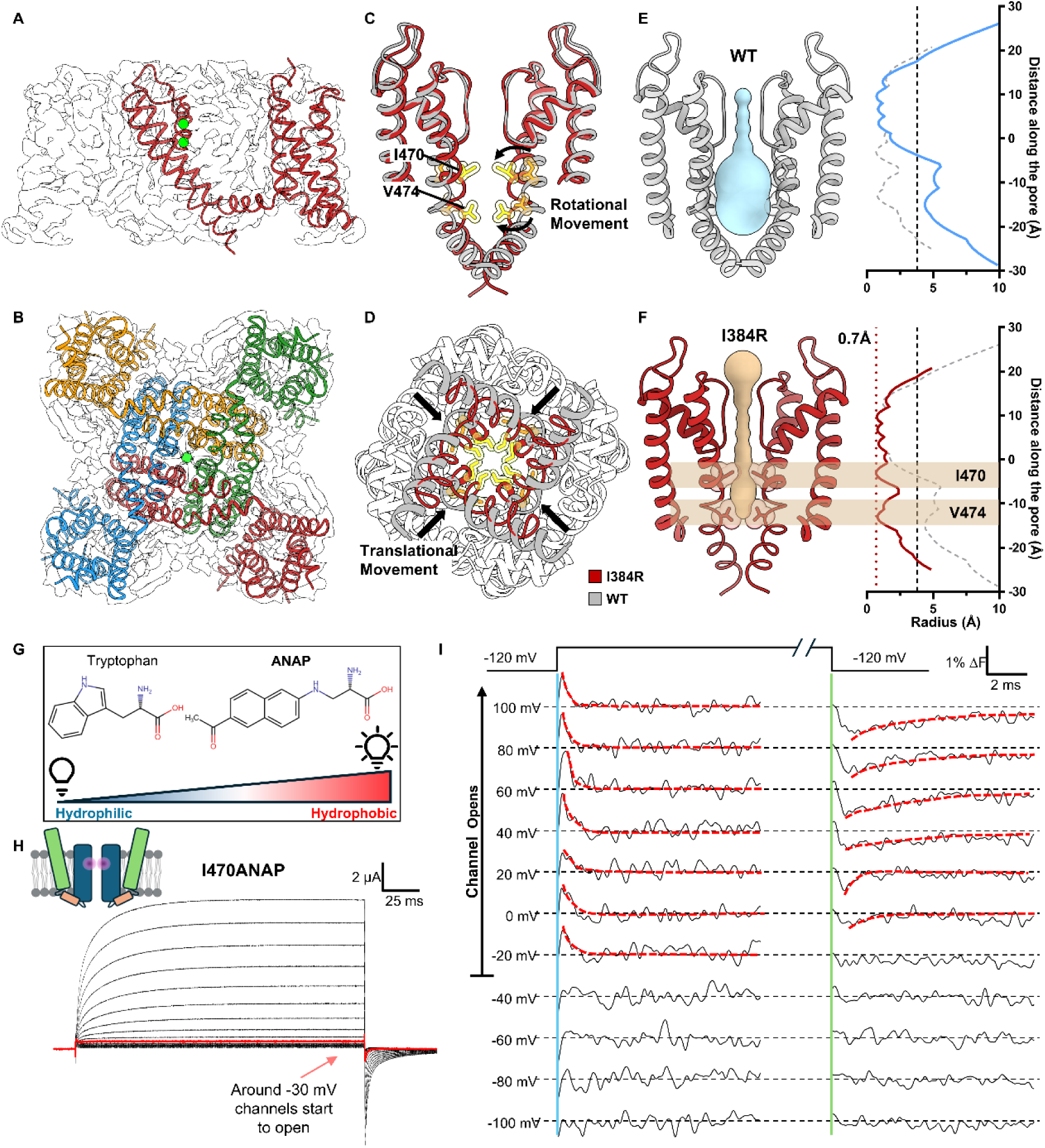
Cryo-EM structure of the completely uncoupled I384R channel, captured in a closed state **A)** Side view of I384R structure. The model is shown overlapping with the electron density, with the intracellular T1 domain omitted. Global resolution is 3.5 Å, with clear densities for all the transmembrane helices. **B)** Intracellular view of I384R. Like the previous solved structure, the channel adopts a domain swapped arrangement. **C)** Structure of the pore in I384R (red) compared to the open state (gray, PDB:7SIP) from the side view. A rotational movement at the upper part of the S6 helices (highlighted by arrows) posits two hydrophobic residues, I470 and V474 from the side direct into the permeation pathway. **D)** Intracellular view of the pore in I384R (red) and WT (gray). In addition to the rotational movement at the extracellular end of the S6 helices, a translational movement towards the pore is observed in the closed state structure, further constraining the pore. **E)** the radius profile of open state pore from the WT channel. The inner cavity and bundle crossing are both open. Dashed vertical line indicates the approximate size of hydrated K^+^ ion. **F)** Radius profile of pore in the uncoupled channel. The pore shows two constriction sites, one at I470 in the inner cavity and the other at V474 in the conserved “PVP” motif. The Narrowest point of the pore is less than 0.7 Å, indicated by the dashed red line. Clearly the pore is captured in the closed state. **G)** ANAP is a fluorescent unnatural amino acid that is sensitive to hydrophobicity of its local environment. With the filter set used in this work, more hydrophobic environment would lead to an increase in fluorescence signal. **H)** Representative ionic traces from I470ANAP. ANAP is incorporated at the 470 position in a site-specific manner through amber stop codon suppression. Robust ionic current could be recorded from I470ANAP. **I)** Fluorescence signal from I470ANAP. A transient signal could be seen among voltages where the channel opens (above -40 mV). A slightly slower signal is seen at the repolarizing pulse as well (highlighted by red dashed line). At more negative voltages however, no such signal could be resolved. It seems the observed fluorescence signal is associated with the opening and closing of the channel.

To validate our structural observations, we set out to measure the local conformational changes around the position 470 in the WT channel utilizing a fluorescent unnatural amino acid probe, ANAP. ANAP has a size comparable to tryptophan which allows minimal disruption of channel function upon incorporation and its fluorescence changes according to the hydrophobicity of its local environment (Fig. 2G) (Chatterjee et al., 2013; Hyun et al., 2009). Utilizing the amber stop codon suppression method (Kalstrup & Blunck, 2017), we incorporated ANAP in a site-specific manner at the 470 position and recorded the ionic current and fluorescence signal simultaneously from I470ANAP (Fig. 2H, I). As channels opened, a fast transient fluorescence change was observed at the start of the depolarizing pulse and a slower one at the beginning of the hyperpolarizing pulse (Fig. 2I, highlighted with dashed line). Clearly, a structural rearrangement occurred at the I470 position as the channels opened and closed. Additionally, the transient nature of the fluorescence signal seems to suggest a rotational movement where the different environments are sampled before reaching its final state, consistent with our structural observations. In the negative voltage range (<-40 mV), where the channel does not open, no fast fluorescent signal was observed, demonstrating that the fluorescence signal at I470 is only observed when the channel opens. These results are fully consistent with the idea that the structure of the uncoupled channel most likely represents a true closed state of the pore and suggest interactions between the bundle crossing region and the selectivity filter.

### A tripartite interaction pocket essential for Electromechanical Coupling

Globally, I384R mutation did not cause a kink in the S4-S5 linker or a local movement at the “elbow” region as was seen before in the resting bacteria Nav channels structures (Wisedchaisri et al., 2019). Instead, we see a lateral, translational movement along the whole length of the S4-S5 linker towards the pore (Fig. 3A). Closer inspection revealed a tripartite interaction pocket among the N-terminus end of the S4-S5 linker, S6 helix in the same subunit and the C-terminus end the S4-S5 linker helix from the adjacent subunit (Fig. 3B). In the strictly coupled WT structure, I384 was securely lodged in a hydrophobic color). The hydroxyl group of Y485, on the other hand, interacted intimately with R394 and E395 in the S4-S5 linker from the adjacent subunit, establishing a structural coupling between the pore and the S4-S5 linker (Fig. 3C – gray color). In the uncoupled I384R structure, however, these interactions are all abolished. The full positive charge of the introduced arginine at position 384 forced itself out of the hydrophobic pocket where the side chain swings towards the adjacent S4-S5 linker. This conformational rearrangement pushes the adjacent R394, facing towards the pore previously, away from the S6 helix and allows for the formation of a salt bridge between R384 and E395 (Fig. 3B, C – red color). These newly formed interactions allowed a closer interaction among the S4-S5 linkers, creating a tight collar around the S5 and S6 segments stabilizing the closed state, and abolished the previous interactions with Y485 in the S6 helices. The observered disruption of the tripartite interactions likely underlies the structural basis for the uncoupling mechanism of I384R, similar to what was shown physiologically elsewhere (Batulan et al., 2010; Chowdhury et al., 2014).

**Figure 3.**
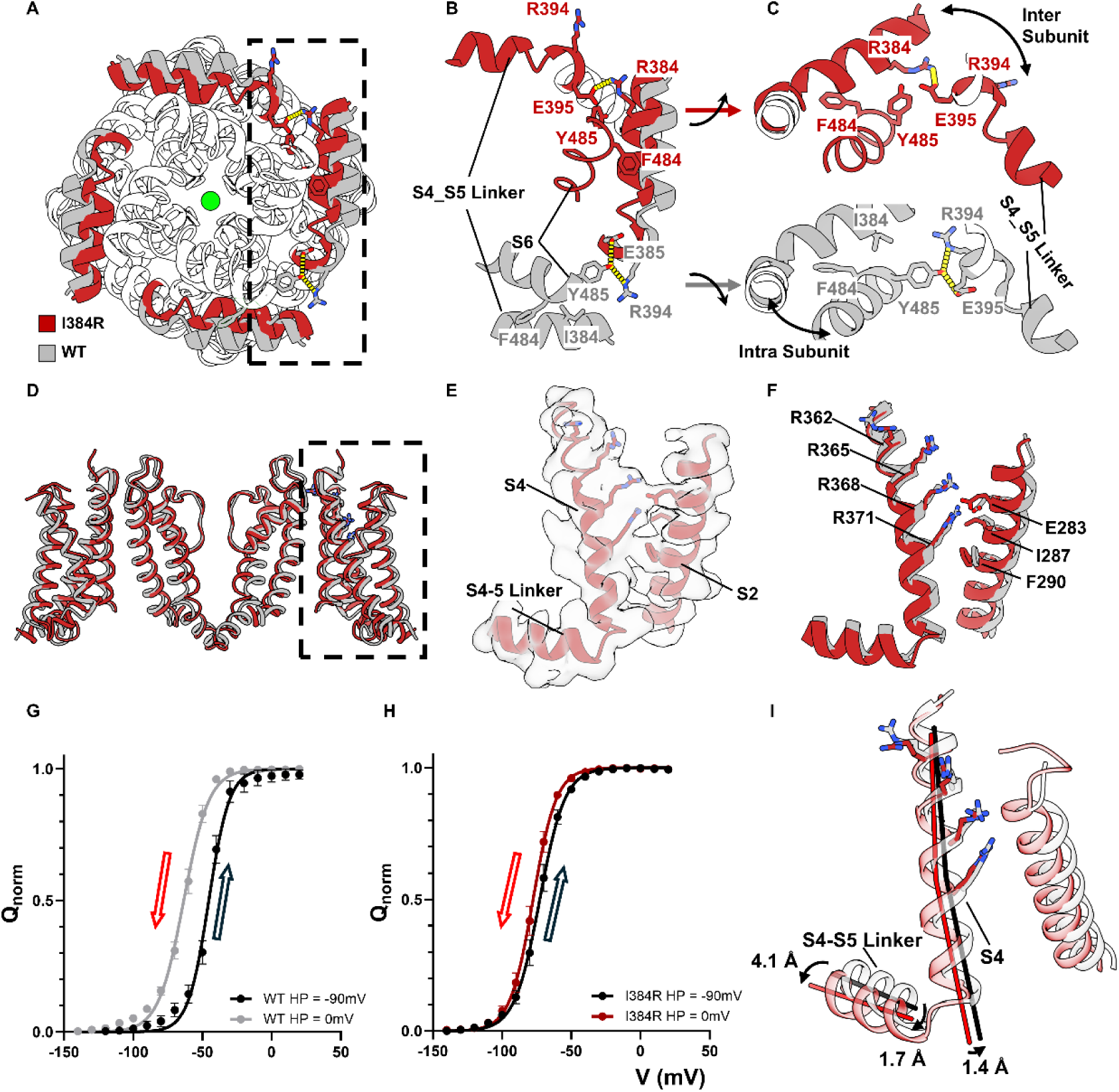
A tripartite interaction pocket for electromechanical coupling and the structure of fully activated but not relaxed voltage sensors. **A)** Intracellular view of the uncoupled structure (red) and coupled structure (gray) with their S4-S5 linkers highlighted in colors. In the uncoupled structure, S4-S5 linkers undergo a translational movement towards the pore, forming a much tighter collar around the pore-forming S6 helices. **B), C)** In the coupled structure (gray), Y445 at the bottom of the S6 helix interacts with E395 and R394 in the C-terminus end of the S4-S5 linker from the adjacent subunit. I384 from the same subunit is lodged in a hydrophobic pocket formed by F484 and Y485. In the uncoupled structure (red), however, R384 jumps out of the hydrophobic pocket and forms a direct salt bridge with E395 in the adjacent unit, which also repulses the R394 away from facing the S6 helix, abolishing completely the interactions with Y445. **D)** Side view of the VSD of the uncoupled channel (red) and the WT channel (gray). **E)** Electron density for the VSD in I384R. All the side chain of the gating charges can be clearly resolved. **F)** Comparison of the WT and I384R voltage sensors. In both cases, all for gating charges (R362, R365, R368 and R371) have moved passed the hydrophobic plug around I287 and F290 region. Both VSDs are in the fully activated state. **G), H)** Hysteresis of the VSDs in WT and I384R, respectively. Holding at 0mV for prolonged time (30s) leads to a ∼20mV shift of the QV curve to the left in the WT channels. However, in the uncoupled channel I384R, no such dramatic shift was observed. It seems at 0 mV, the VSDs in I384R does not enter into the relaxed state. **I)** Helical movements in S4 due to the S4-S5 linker. The tight conformation of the S4-S5 linker shifted the C terminus of the S4-S5 linker for 4.1 Å. This shift is transduced to the N terminus end of the linker and the S4 helix as well, shifting them 1.7 Å and 1.4 Å, respectively.

### Voltage-sensing domain captured in fully up but not relaxed state

Gating current measurements demonstrated that at 0mV, the QV curve of I384R had reached its maximum, suggesting that all the voltage sensors had activated. However, since there are multiple intermediate states for the VSDs, the electrophysiological data cannot unequivocally define whether the voltage sensors reach the fully activated state in the uncoupled channel or even if they move in a similar way as the WT channels. To address this, we compared the VSD structures in I384R and the WT (Fig. 3D). Clear densities were resolved for all gating charges in S4 (R362, R365, R368, R371) (Aggarwal & MacKinnon, 1996; Seoh et al., 1996) as well as the key residues that form the hydrophobic plug in S2 (I287, F290) (Lacroix et al., 2014; Lacroix & Bezanilla, 2011; Tao et al., 2010) and the countercharge (E283) (Fig. 3E). Our structure shows that all four gating charges have moved pass the hydrophobic plug in the uncoupled conformation and are accessible to the extracellular solution (Fig. 3F) similar to the case for the WT structure. Clearly in the uncoupled channel, the voltage sensors moved similarly to the WT channels and reached the fully activated state.

However, unlike the WT, the voltage sensors in I384R do not appear to enter the relaxed state, a conformation that has been observed in Kv, Nav, Cav channels and voltage sensitive phosphatases (VSP) and is driven by prolonged depolarization (Bezanilla, 2018; Bezanilla et al., 1982; Lacroix et al., 2011; Olcese et al., 1997; Shirokov et al., 1992; Villalba-Galea et al., 2008). In WT channels with W434F background, holding the channels at 0 mV shifts the QV curve almost 20 mV to more negative potentials when compared to holding at -90 mV, a hallmark of entry into the relaxed state (Fig. 3G). This was not observed in the uncoupled channel. Holding I384R at 0mV for a long period of time (> 30s) did not cause significant shifts in the QV curve, showing that the voltage sensors in I384R did not enter in the relaxed state, at least at this voltage (Fig. 3H). Since the cryo-EM structures were captured at 0 mV, the physiological evidence would then argue that the structure of VSDs in the open channel represented the relaxed state while in the uncoupled channel, the VSDs likely resided in the non-relaxed state. Structurally speaking, the major difference between VSDs in the WT and I384R lies mostly in the displacement of the alpha helixes. In I384R, the S4-S5 linkers were more intimately interacting with each other, resulting a big lateral shift particularly at the C-terminus end of the helix, 4.1 Å away from the WT structure (Fig. 3I). This movement was transduced to the N-terminus end of S4-S5 linker and the S4 helixes, dragging them 1.7 Å and 1.4 Å away from the open state structure, respectively. Since the entry of the relaxed state has been associated with opening the pore (Haddad & Blunck, 2011), it is possible that the observed helical displacement in I384 formed the structural basis for the relaxed state of the VSDs.

### Noncanonical conformation of the selectivity filter and the decreased volume of the closed state channel

One surprising observation of the I384R closed state structure comes from the selectivity filter conformation. Instead of the typical 4 K^+^ ion densities seen in the conductive filter (Fig. 4B) (Long et al., 2007; Tan et al., 2022), only 2 putative bound K^+^ ions at S2 and S4 sites within the SF can be resolved (Fig. 4A). It is intriguing to see that the S3 K^+^ was absent in the structure, since generally, the S3 position typically displays the strongest K^+^ occupancy in K channel structural determinations (Lee et al., 2025). In our case, the coulombic density suggests that occupancy was similar for these two positions with a slightly higher occupancy at S4 position (Supp. Fig. 6). Structurally, two small twists were observed at the S1 and S4 binding site when compared to the WT structure (Fig. 4C). The carbonyl group of the last glycine in the TVGYG selectivity filter, G446, flipped away from the pore, directly altering the binding site at S1 position (Fig. 4D). A similar twist was also observed at the bottom of the selectivity filter at the T442 position, which might account for the slight shift of the K^+^ ion at the S4 position compared to the WT.

**Figure 4.**
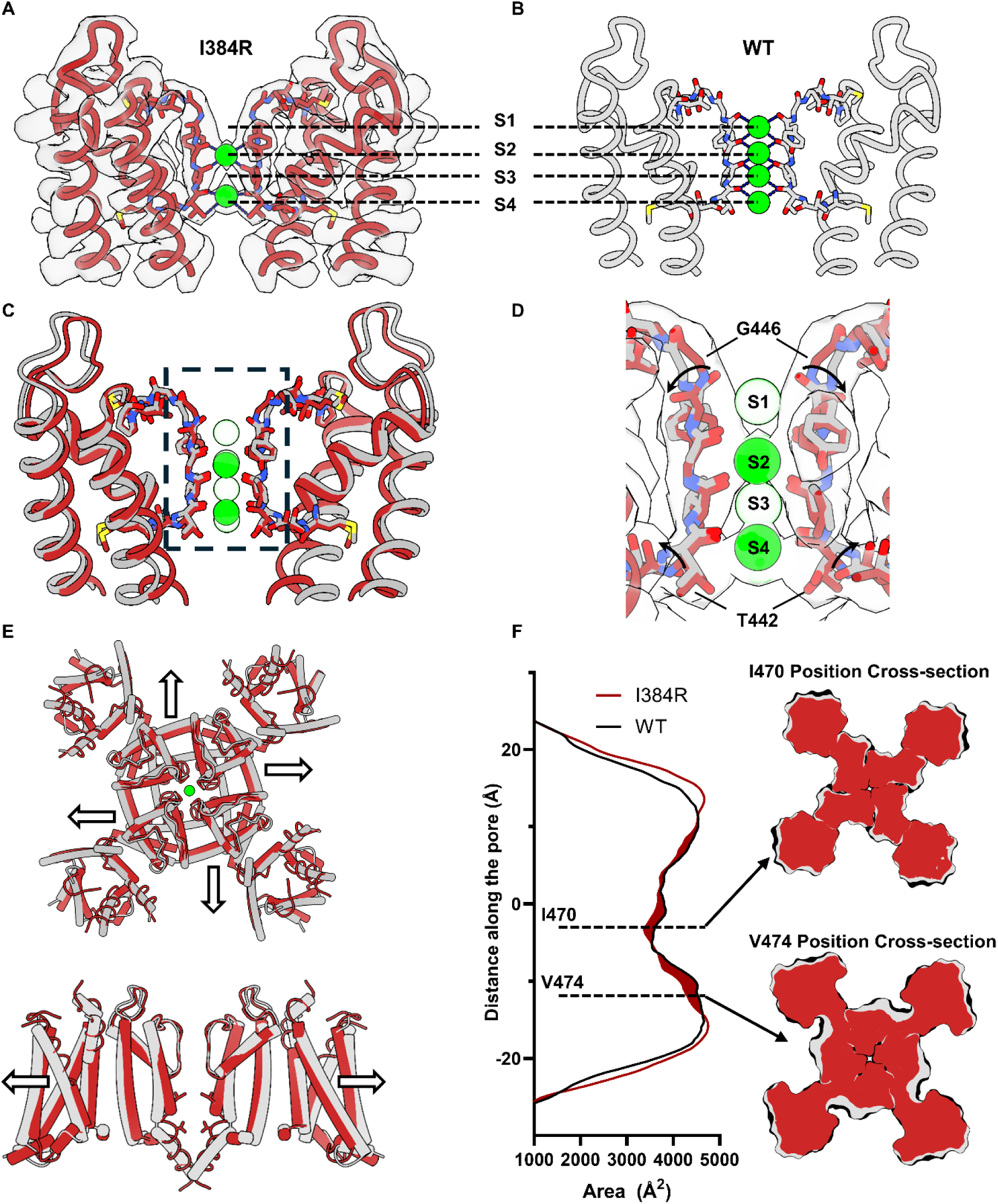
Noncanonical conformation of the selectivity filter and global decrease in protein volume in the closed state. **A)** The SF captured in closed state. The atomic model is overlaid with the density map. Clearly there only exist two K^+^ in the SF. **B)** Conductive SF captured in the open state. (PDB:7SIP). **C)** Overlay of the SF in the closed (red) and open state (gray). In the closed state, K^+^ ions are seen bound at S2 and S4 position. **D)** Difference in SF between open and closed state structure. The carbonyl on the backbone of G446 flipped away from the permeation pathway, abolishing the S1 binding site on the top of the selectivity filter. A similar twist is seen at the bottom of the SF at T442 position, resulting in small displacement of the threonine side chain. **E)** The channel is more expanded in the open state (gray) compared to the closed state (red). The expansion can be seen in the VSDs as well as the pore. **F)** Cross section area calculation utilizing CHARMM_GUI.

Another intriguing observation in the closed state structure was the decrease of the protein volume in the transmembrane region. When comparing the WT with I384R structures, we noticed that the protein expanded laterally in the open state (Fig. 4E). This expansion was due to the translational movement of S4-S5 linker and the S6 helices. Area calculations with CHARMM_GUI show an asymmetric increase of the cross-section area of the open channel compared to the closed one (Fig. 4F) (Jo et al., 2008). The most significant expansion happened around the I470 and V474 region, where the cross section of the open channel increased by almost 10% (Supp. Fig. 7). This expansion in volume might be the underlying mechanism of the reported mechanosensitivity of the Kv1 channels (Laitko & Morris, 2004; Tabarean et al., 1999).

## Discussion

### Electromechanical coupling in voltage-gated ion channels and its energetics

All voltage-gated channels share two basic functional modules: the voltage sensing domain and the pore domain. Electromechanical coupling describes the communication between these two modules. In the present study, we identified a tripartite pocket that is essential for electromechanical coupling in the Shaker potassium channel, a strictly coupled channel. Structurally, interactions among Y485, F484, I384, E395 and R394 establish the functional connectivity between the pore and the S4-S5 linker. These intersubunit interactions likely contribute to the cooperativity of the voltage sensors and the pore opening as well (Zagotta et al., 1994). It has been demonstrated previously that reducing the side chain volume at positions Y485 and F484 in the S6 leads to shallower and right-shifted GV curves, (Ding & Horn, 2003; Hackos et al., 2002; Haddad & Blunck, 2011), similar to what we show here in I384E, I384L and I384N. Mutagenesis experiments and thermodynamic cycle analysis among E395, R394 and Y485 have confirmed their energetic coupling and demonstrated their importance for the electromechanical coupling (Batulan et al., 2010; Chowdhury et al., 2014; Yifrach & Mackinnon, 2002). The identified tripartite pocket is thus likely responsible for the efficient transfer of the movement from the voltage sensor to the pore seen in the Kv channels.

In strictly coupled voltage-gated channels, the opening of the pore requires obligatory coupling of the voltage sensors and pore. Understanding the energetics of this coupling is of fundamental importance to define the nature of EMC. It has long been debated whether it is energetically favorable for the pore to stay in the open or the closed state. In other words, are the voltage sensors doing work to “pull” the channel open or to “push” to keep the pore closed. While some computational work suggests the pore prefers to stay open in the absence of an external energy bias (Fowler & Sansom, 2013), our physiological results seem to argue otherwise. In the I384 mutants, all the uncoupled mutants (I384L, I384E and I384N) show a right shifted GV curve and a left shifted Q-V curve, as if the pore load to the sensor is decreased. In these partially uncoupled mutants these results indicate that the pore is now less firmly attached to the VSD, making it more difficult to open, preferring to stay in the closed state.

### Kv channel structure captured with a closed pore

Among the uncoupling mutants studied, I384R is the most extreme case where no ionic current is recorded, and only gating current is seen. This is quite different from the classical W434F mutant, even though they both have minimal ionic conductance. W434F mutant speeds up the C-type inactivation and stabilized the channel in the inactivated state (Yang et al., 1997). Thus, I384R represented an ideal candidate to probe the conformation of the inner bundle gate, and we were able to capture the closed pore structure of Shaker potassium channel by disrupting the electromechanical coupling (Fig. 1, 2). It is worth pointing out that this uncoupled pore most likely represented the closed state, instead of an inactivated state, another nonconductive state in Kv channels. It is the EMC that was altered by substituting isoleucine at position 384, as indicated by the relative shifts in QV and GV curves (Fig. 1 & Supp Fig. 1). There are informative differences in the behavior of the gating currents measured in mutants W434F and I384R. In I384R, the gating current was significantly faster. By severing the electromechanical coupling between PD and VSD, the sensor is free to move in the absence of a mechanical “load”: we are not tampering with the PD itself and therefore, the structure of the closed pore likely represents the true closed state of the pore in the WT channels. In contrast, gating currents in W434F, are a reflection of the VSD movement under a physiological load. The suppression of ionic currents is derived from effects downstream to the activation gating.

A surprising finding from the closed pore structure is the large degree of conformational changes happening along the entire length of the S6 helices. In contrast to the simple hinge-like movement seen in prokaryotes (Cuello et al., 2017; Tombola et al., 2006). Inner gate opening in eukaryotic strictly coupled Kv channels appears to be more complex, with both, a lateral movement of the PVP motif plus an additional rotational movement at the inner cavity of pore. This type of movement leads to the collapse of the inner cavity in the closed state, where hydrophobic I470 rotates and points directly into the permeation pathway. Our fluorescence experiments with ANAP directly demonstrated that a conformational change happens near position 470 as the channels open and close, studies with tetraethyl ammonium (TEA) ion. TEA exerts its blocking effects by entering the inner cavity of the channel in the open state and needs to be expelled out the channel before the pore closure can happen, a foot in the door effect (Armstrong, 1971). Mutating I470 to a smaller residue like cysteine, allows TEA to stay in the inner cavity in the closed state (Holmgren et al., 1997).

### Conformation of selectivity filter in closed and open state

The selectivity filter in the I384R closed state structure was captured in what appears to be a noncanonical conformation. Current SF conformation is different from the typical conductive filter where four bound K^+^ ions orderly occupy the S1-S4 positions. It is also dissimilar to alternative “dilated” inactivated filter where the dilation of the top part of the filter abolish S1 and S2 binding sites (Supp. Fig. 8A) (Stix et al., 2023) or the “pinched” inactivated filtered captured in prokaryotic Kv channel (Supp. Fig. 8B). In I384R, two bound K^+^ ions were found at S2 and S4 position in the absence of any expansion at Y445 position or pinching at G444 (Cuello et al., 2010) (Supp. Fig. 8). It is then possible that as the channel opens, a conformational change around the SF occurs. Some existing fluorescence measurements seem to be in line with this hypothesis; TMR labeled channels at T449 position, around the selectivity filter region, gave rise to a fluorescence change when the channels opened and the FV curve followed very closed the GV curve (Cha & Bezanilla, 1997). In Nav channel, similar phenomenon has been suggested in the fast inactivated state as well (Liu et al., 2023). It is possible that the S6 movements during activation are allosterically communicated to the selectivity filter, triggering the transition from the noncanonical conformation to the typical conductive conformation (Peters et al., 2013). However, further investigation is certainly required before a definitive conclusion can be reached.

### Isoleucine 470 as a shared structural element for both activation and inactivation

Pore radius calculation identified I470 in the inner cavity as the upper gate in closed state structure. This is a very interesting observation because I470 has been mostly associated with the C-type inactivation of Kv channels, previously. Physiological experiments mutating I470, together with T449, demonstrated that the inactivated state was rendered conductive by the double mutations (Olcese et al., 2001), and computationally, long-time scale simulation also observed a coincidence of I470 movement and closure of the permeation pathway in the inactivated channels (Treptow et al., 2024). Moreover, structural results from Kv4.2 identified the role of the equivalent isoleucine in inactivation mechanisms (Ye et al., 2022). In the case of Shaker K^+^ channel, the inactivation state is entered after the activation of the channel and the open inactivated state is the most dominant inactivated state in the channel (Panyi & Deutsch, 2006, 2007; Szanto et al., 2020). It appears that I470 is a shared structural element for both activation and inactivation and could serve as the link for the interplay of these two functional states.

### Mechanosensitivity of Kv channel gating

In the closed state structure, we saw a collapsed inner cavity in the center of the protein which led to a global shrinkage of the protein volume. Compared to the open state structure, the protein occupied less area on the membrane in the closed state. It is worth noting that the open state structure was solved in lipid nanodisc and the current closed state structure was solved in detergents. The difference in structural determination method may result in an underestimation of the actual change in protein volume between closed and open state since it has been reported that reconstitution of proteins into nanodisc could introduce additional lateral forces from the lipid bilayer to the proteins (Dalal et al., 2024). Therefore, it is possible that the actual cross section area of the channel in the closed state would be smaller and the actual expansion from closed to open would be more significant in a physiological condition.

One prediction coming from the observed decrease of the protein volume is that to open the channel, additional energy is required to push against the lateral pressure from the membrane and the amount of the lateral pressure will modulate the gating of the channel, giving rise to some mechanosensitivity to the channel. Some physiological experiments support this prediction. Excised patch clamp experiments have demonstrated that the GV curve of Shaker-IR-WT channels shifted to the left, opening more easily, with increased suction in the pipette (Laitko & Morris, 2004; Morris, 2011; Schmidt et al., 2012). Additionally, mechanical modulation on the gating of the channel seems to be mostly limited to the change in the V_1/2_, which mostly reflects changes in steady-state energy level, without altering the number of charges that move. It is possible that the transition from the closed to the open state requires a global expansion of the protein which could serve as the molecular basis of the mechanosensitivity in strictly coupled Kv channels and possible other members of voltage gated channels, such as Nav channels (Morris & Juranka, 2007).

### Activation mechanism for strictly gated Kv channels

Current experimental results argue for a new gating model for strictly gated Kv channels (Fig. 5). At very hyperpolarizing voltage, the channels occupy the deepest closed state, where all four VSDs are in the down state. In this deep closed state, the permeation pathway is gated by I470 in the inner cavity as well as V474 at the bundle crossing region. Upon depolarization, asynchronous activation of the VSDs moves the channels into an ensemble of intermediate closed states, where some but not all VSDs are activated. The movement of the S4 in the VSDs creates a direct pull on the S4-S5 linker. However, this pull does not provide enough energy to open the channel and the VSDs are stabilized in the non-relaxed state. When all four voltage sensors move up, the channels enter a transient pre-open state, which is represented by the structure of the uncoupled channel.

**Figure 5.**
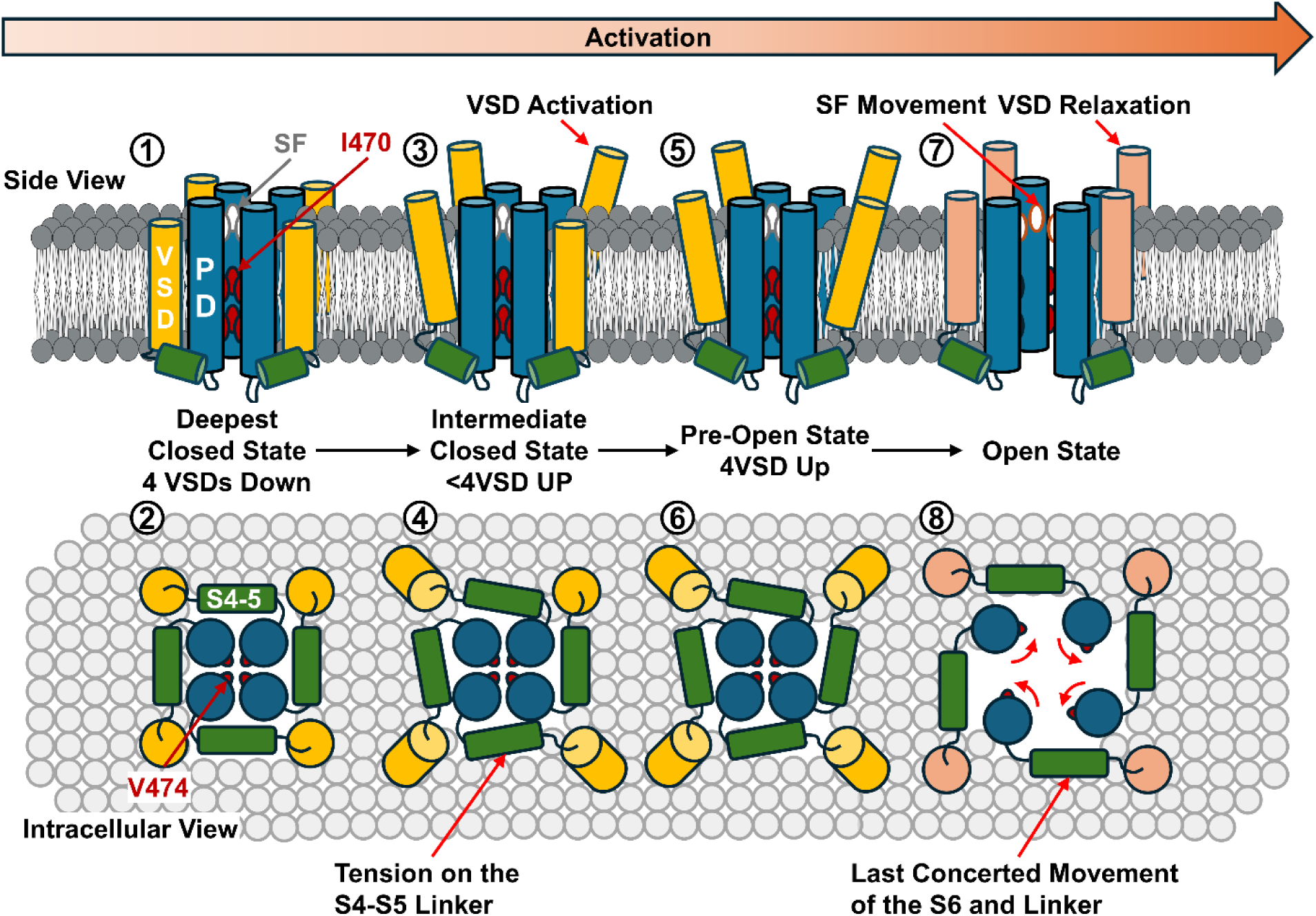
Proposed activation mechanism and interactions with different functional states in Kv1 channels. The progression to activation is from left to right. In the deepest closed state of the channel, all the VSDs are down, and the SF likely resides in the noncanonical state (①). The closed pore is sealed closed by hydrophobic I470 and V474 (①, ②). Upon depolarization, VSDs transit to active but not-relaxed state (③). The upward movement of the VSDs creates a pull on the S4-S5 linker, yet since not all VSDs are up, the energetic input is not enough to open the pore (④). In the last closed state before opening, or the pre-open state, all four VSDs move up (⑤), creating enough pull on the S4-S5 linkers (⑥). In the case of I384R, the channel is most likely stabilized in this pre-open state with newly introduced salt bridges. From the pre-open state, the last concerted movement happens and the S6 helices undergo a roll and turn movement, opening up the permeation path and expanding the channel laterally (⑧). At the same time as the pore opens, the lateral movement of the S4-S5 linker drives the VSDs into a different state, the relaxed state (⑦).

In this pre-open state, all the VSDs populate the activated (up) conformation which provides enough energy into the linkers to drive the pore into its open conformation. As part of a last concerted movement. S4-S5 linkers expand, allowing the opening of the pore and the relaxation of the VSDs. During opening, the S6 undergoes a roll and turn movement that rotates away the upper and lower activation gate, creating a water-filled permeation pathway. This movement is allosterically communicated to the selectivity filter leading to a small local conformation change, allowing the selective permeation of the K^+^ ions and predisposing itself for C-type inactivation, forming the structural and molecular basis for interaction among different functional states.

## Materials and Method

### Site-directed mutagenesis and cRNA *in-vitro* synthesis

Shaker zH4 K+ channel with fast inactivation removed (Δ6–46 - Shaker-IR (Hoshi et al., 1990))in pBSTA vector, flanked by β-globin sequences, was used in this study for all the physiological experiments. Point mutations were generated using mismatched mutagenic primers. The PCR product was digested in DpnI to remove the template and was used to transform the XL-gold ultra-competent cells. After ampicillin resistance screening, plasmids were purified from the colonies using standard miniprep protocols. Purified plasmids were sent for either Sanger (Genomics Facility at University of Chicago) or nanopore (Plasmidsaurus) sequencing to confirm the introduction of the point mutation and the absence of off-target mutations. cRNA was then transcribed *in vitro* from the linearized plasmids (T7 RNA expression kit; Ambion Invitrogen, Thermo Fisher Scientific, Waltham, MA).

### *Xenopus laevis* oocyte preparation and channel expression

Ovaries of *Xenopus laevis* were purchased from XENOPUS1 (Dexter, Michigan). The follicular membrane was removed using collagenase type II (Worthington Biochemical Corporation) 2 mg/mL with bovine serum albumin at 1mg/mL (BSA). After defolliculation, stage V–VI oocytes were then selected and microinjected with 5-100 ng cRNA. Injected oocytes were incubated at 18 °C for 1-5 days in SOS solution (in mM: 100 NaCl, 5 KCl, 2 CaCl_2_, 0.1 EDTA, and 10 HEPES at pH 7.4) supplemented with 50 µg/mL. Unless otherwise stated, all chemicals were purchased from Sigma-Aldrich.

### Cut-open voltage clamp on *Xenopus laevis* oocytes

Macroscopic ionic and gating current were recorded using cut-open voltage clamp technique (Stefani & Bezanilla, 1998). Micropipettes filled with 3M CsCl or NaCl, with resistance between 0.4 and 0.8 MΩ were used to measure the internal voltage of the oocytes. Current data were online filtered at 20 kHz with a low-pass 4 pole Bessel filter and sampled by a 16-bit A/D (USB-1604; Measurement Computing, Norton, MA) converter at 1 MHz. All experiments were conducted at room temperature (∼ 17 °C). For ionic current experiments, unless otherwise stated, were conducted in external solution consisted of in mM: 12 K methylsulfonate (MES), 108 N-methyl-D-glucamine (NMG) MES, 2 Ca MES, 10 HEPES, 0.1 EDTA, pH = 7.4 and internal solution consisted of in mM: 120 K MES, 10 HEPES, 2 EGTA, pH = 7.4. The capacitive transient was manually compensated with a dedicated circuit and in some cases, further removed by an online P/-4 protocol with a holding voltage of either -80 or -90 mV (Armstrong & Bezanilla, 1977).

For gating current experiments, all experiments were conducted in external solutions consisted of in mM: 120 NMG MES, 2 Ca MES, 10 HEPES, 0.1 EDTA, pH = 7.4 and in internal solution consisted of in mM: 120 NMG MES, 2 Ca MES, 10 HEPES, 2 EGTA, pH = 7.4. The gating current was recorded with W434F background with the exception of I384R, which by itself produced no discernable ionic current.

### Single channel recordings and noise analysis

Single channel recordings and noise analysis were performed on excised inside-out *Xenopus Laevis* oocyte patches. Briefly, the cells were injected with 5ng of RNA and maintained at 12 C° the day before the experiment. To remove the vitelline membrane, oocytes were incubated for 5 minutes in a hypertonic solution (SOS solution supplemented with 300 mM sucrose). This procedure shrank the oocyte separating the plasma membrane from the vitelline membrane, making the mechanical removal of the vitelline membrane easier without compromising the integrity of the cell. After removal of the vitelline membrane, oocytes were immediately and gently washed three times with intracellular solution containing in mM: 120KMES, 2EGTA, 10HEPES, pH = 7.40. They were placed in a recording chamber onto an inverted microscope. Current was recorded with an Axopatch 200B patch-clamp amplifier. The pipette solution consisted of, in mM, 120KMES, 2KCl, 2CaMES, 10HEPES, 0.1 EDTA, pH = 7.40. The resistances of the tips were between 13∼17 MΩ for single channel experiments and 6∼10 MΩ for noise analysis experiment. To reduce stray capacitance, the tips of the pipette were covered by Sylgard 184 (Dow Corning Corporation). Current was filtered with a digital 8 pole Bessel filter set at 10kHz (3384 Krohn-Hlite). For noise analysis, we applied hundreds of depolarizing voltage pulses. The ionic currents elicited were averaged to obtain the mean. The variance and the mean were obtained using our Analysis software. For single channels, we recorded several current traces from voltage pulses to +140mV. The histograms were obtained using Analysis.

### Unnatural amino acid incorporation and voltage-clamp fluorimetry

To incorporate ANAP, we utilized the amber suppression technique. Briefly, the oocytes were injected with a mixture of ANAP-synthetase, ANAP methylester, ANAP-tRNA, *Xenopus* release factor 1 with D55E mutation and messenger RNA encoding the channel with the amber stop codon introduced at I470 position (Chatterjee et al., 2013; Durner et al., 2023; Hyun et al., 2009). The oocytes were incubated in the dark for 3 to 5 days prior to recording. The voltage clamp fluorometry setup and the filter set used in the study was similar to what was previously described (Kalstrup & Blunck, 2013, 2017). However, instead of using a photo diode, a photomultiplier was used to maximize the sensitivity of the signal, while allowing a decrease of the excitation light, decreasing photobleaching. The fluorophore was excited with a LED at 365nm (Thor lab M365L3). TMR labeling was done similarly as previous described (Cha & Bezanilla, 1997). Briefly, oocytes were incubated in 1mM DTT for 15 minutes. Three washes were administered before transferring the cells to the labeling solution with 20 µM Tetramethylrhodamine-5-Maleimide (Thermo Fisher T6027) in a depolarizing solution (in Mm: 120 KMES, 2 CaMes, 10HEPES, 0.1EDTA, pH = 7.40) for 30 minutes.

### Data Analysis

Ionic current was taken by the steady-state current level and converted to conductance using the following relationship:

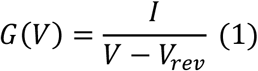

where*, I* is the ionic current in steady state, *V* is the membrane voltage, and *V_rev_* is the reversal potential for the conducting ion. The GV curves were then fitted using a two-state model given by the equation:

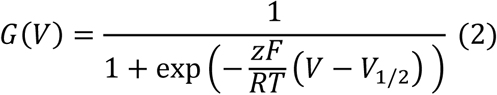

where *z* is the apparent charge expressed in units of elementary charge (*e*_0_), *V* is the voltage and *V_1/2_* is the voltage of half-maximal conductance. *R* is the ideal gas constant, *T* is the temperature in Kelvin and *F* is the Faraday constant.

The gating charge was obtained by integrating the on and off-gating currents. They were plotted against the voltage to obtain the QV curves.

For the analysis of the normalized QV curves, we used a two-state model fitting equivalent to the one in Equation 2 to fit the gating from the I384E, I384L, I384N, and I384R mutants. The equation was the following:

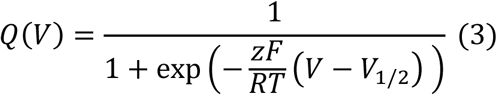

For WT, I384A, I384C, we used a three-state model fitting given by the following equation:

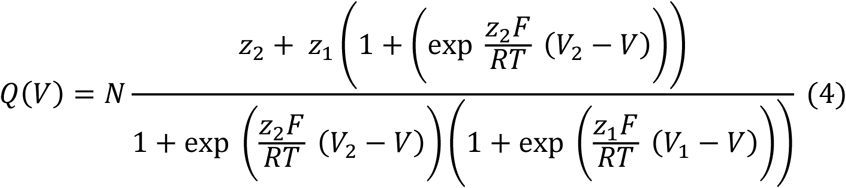

where *N, z_1_, z_2_, V_1_ and V_2_* are the number of channels, the charges associated and equilibrium voltages for the first and second transition, respectively.

For nonstationary noise analysis, the mean variance data were fitted using a parabolic equation as follows:

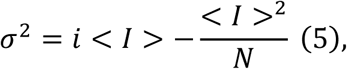

where σ^2^ is the variance, *i* is the single-channel current, < *I* > is the mean current, and *N* is the number of channels in the patch. We can estimate the maximal open probability (*Po*_*max*_) by knowing the maximum mean current (*I*_*max*_) using the following equation:

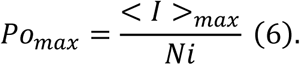

For single channel analysis data, we obtained all-points histogram binned at 0.05 pA using our Analysis software. The data was then fitted using a mixture of *k* Gaussian Distributions as follows (Colquhoun & Sigworth, 1995):

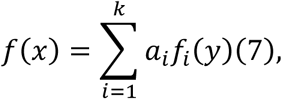

Where *a*_*i*_ and *f*_*i*_(*y*), are, respectively, the relative areas of the components and the Gaussian equation described by:

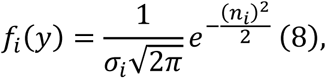

And

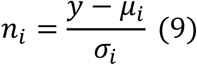

Where μ_*i*_ and σ_*i*_ are the mean and the standard deviation of the individual component *i*.

Data is presented as Mean ± SEM. For noise analysis, ionic and gating currents experiments we used at least 3 oocytes from different batches. For single channels, we recorded several oocytes and only for presentation purposes show the representative traces from one cell.

### Expression and purification of mutant shaker potassium channel

Shaker-IR-I384R was subcloned into a modified pEG BacMam vector containing a C-terminal HRV 3C protease site, an eGFP tag and an 8×-His tag utilizing the 5′ NotI and 3′ XbaI restriction sites. The Bacmid plasmid was generated using the Bac-to-Bac system which was then used to transfect the Sf9 insect cells with Cellfectin (Thermo Fisher). After 4 to 7 days of incubation at 27°C, P_0_ virus was then collected after removing the remaining cell and debris. P_0_ was then amplified to produce P_1_ and P_2_ virus. 2L of HEK293S GnTI^-^ cells were infected with 200mL of P_2_ virus. After 24 h of shaking incubation at 37°C, sodium butyrate was added to the cells at a final concentration of 10mM and the culture was transferred to 30°C. Cells were collected 48 h after infection. The cell pellets were then washed with phosphate-buffered saline (PBS) at pH 7.4, collected by centrifugation, flash-frozen and stored at −80 °C for later purification.

All purification steps were conducted at 4°C. Frozen cell pellets were thawed in water bath and homogenized with a Dounce homogenizer in suspension buffer (150mM KCl, 2mM TCEP, 1mM EDTA, 50mM TRIS, Pierce protease inhibitor tablet (Thermo Fisher Scientific) PH 7.5). Resuspended cells culture were supplemented with DDM:CHS (10:1) at 1% final concentration (m. / v.) and were extracted for 2 hours with gentle agitation. The cell debris was pelleted by ultracentrifugation for 1 hr at 40000 rpm in Ti45 rotor (Beckman Coulter). The supernatant was collected and incubated with 2 ml CNBR-activated Sepharose beads (GE Healthcare) coupled with 4 mg high-affinity GFP nanobodies purified in-house for 3 hours. The Sepharose beads were then rinsed three times 10 column volume with the suspension buffer supplemented with 0.1% DDM:CHS (10:1), 0.05% DDM:CHS + 0.02% GDN, 0.02% GDN respectively. Then beads were incubated with HRV 3C protease overnight to release the purified channels. The next day, 3 column volume of suspension buffer was used to elute the beads. Eluted solution was concentrated with Amicon Ultra Centrifugal Filter unit (Millipore) with 100kDa cutoff to approximately 1mL and loaded onto a Superose6 (10 × 300 mm) gel filtration column (GE Healthcare) and separated with suspension buffer supplemented with 0.02% GDN.

### Cryo-EM sample preparation and data acquisition

Samples purified from size-exclusion chromatography were pooled and concentrated to around 0.7 mg/mL measured with a nanodrop machine. 3.5 µL concentrated protein was applied to glow-discharged (30 s, 20 W Solarus Plasma Cleaner) Quantifoil grids (R 1.2/1.3 Au 300 mesh). The grids were blotted for 3s with blot-force 3 in a FEI Vitrobot Mark IV (Thermo Fisher) chamber with 100% humidity at room temperature before plunge froze in liquid ethane. Clipped grids were subsequently loaded onto a Titan Krios microscope. Single-particle movies were acquired using a K3 direct electron detector in super-resolution mode, coupled with a 20 eV GIF energy filter. Data collection was performed at a nominal magnification of ×81,000, corresponding to a pixel size of 0.5315 Å, binned by 2 during acquisition. The total electron dose was calibrated to 60 e−/Å², distributed across 50 frames per movie.

### Single-particle cryo-EM analysis

All steps for structure determination were performed using CryoSPARC (Punjani et al., 2017), including motion correction and contrast transfer function (CTF) estimation. A subset of 2,000 particles was initially picked and classified in 2D to generate templates for template-based particle picking. Approximately 4,500,000 initial particles were picked and subjected to multiple rounds of 2D classification. From these, 180,000 particles were selected to generate three *ab initio* models with C1 symmetry. Particles from the best class (∼116,000) were then processed for 3D refinement with C4 symmetry, yielding a nominal resolution of 5.7 Å. We performed 3D refinement using non-uniform refinement algorithm (Punjani et al., 2020), which consistently yields best result in our data. A subsequent 3D classification was performed using the best-refined model from the previous round and a junk model generated *ab initio*. The best class from this step was further refined using C4 symmetry, resulting in a 3.55 Å nominal resolution. After identifying a subset of ∼80,000 particles for final refinement, Reference-Based Motion Correction was applied to estimate per-particle movement trajectories and empirical dose weights. A final 3D refinement step was performed, followed by postprocessing using a tighter mask and C4 symmetry enforcement. Local resolution was calculated using CryoSPARC. Validation was done using MOLprobity (Williams et al., 2018).

### Model building and structural refinement

The Shaker-IR model (PDB 7SIP) was used as a template to build atomic models for the Shaker-IR-I384R mutants into our density map. All structural models were constructed using unsharpened maps. Initial models were generated through iterative rounds of manual model building in COOT (Emsley et al., 2010) and real-space refinement in Phenix (Afonine et al., 2018). Final refined atomic models were obtained using interactive flexible fitting in ISOLDE (Croll, 2018). All structural analyses and figure generation were performed using UCSF ChimeraX (Goddard et al., 2018; Pettersen et al., 2021).

## Acknowledgements

We would like to thank Gethiely Gasparini and Hlafira Polishchuk for oocytes preparation, DNA mutation and RNA preparation. We would like to thank Dr. Chris Ahern and Dr. Jason Galpin for helping with the ANAP experiments We would also like to thank the staff at the cryo-EM core in the University of Chicago for their assistance. Supported by NIH grant R01-GM030376, R01-GM150272 and National Science Foundation Award QuBBE QLCI (NSF OMA-2121044).

---

**Supplementary Figure 1.**
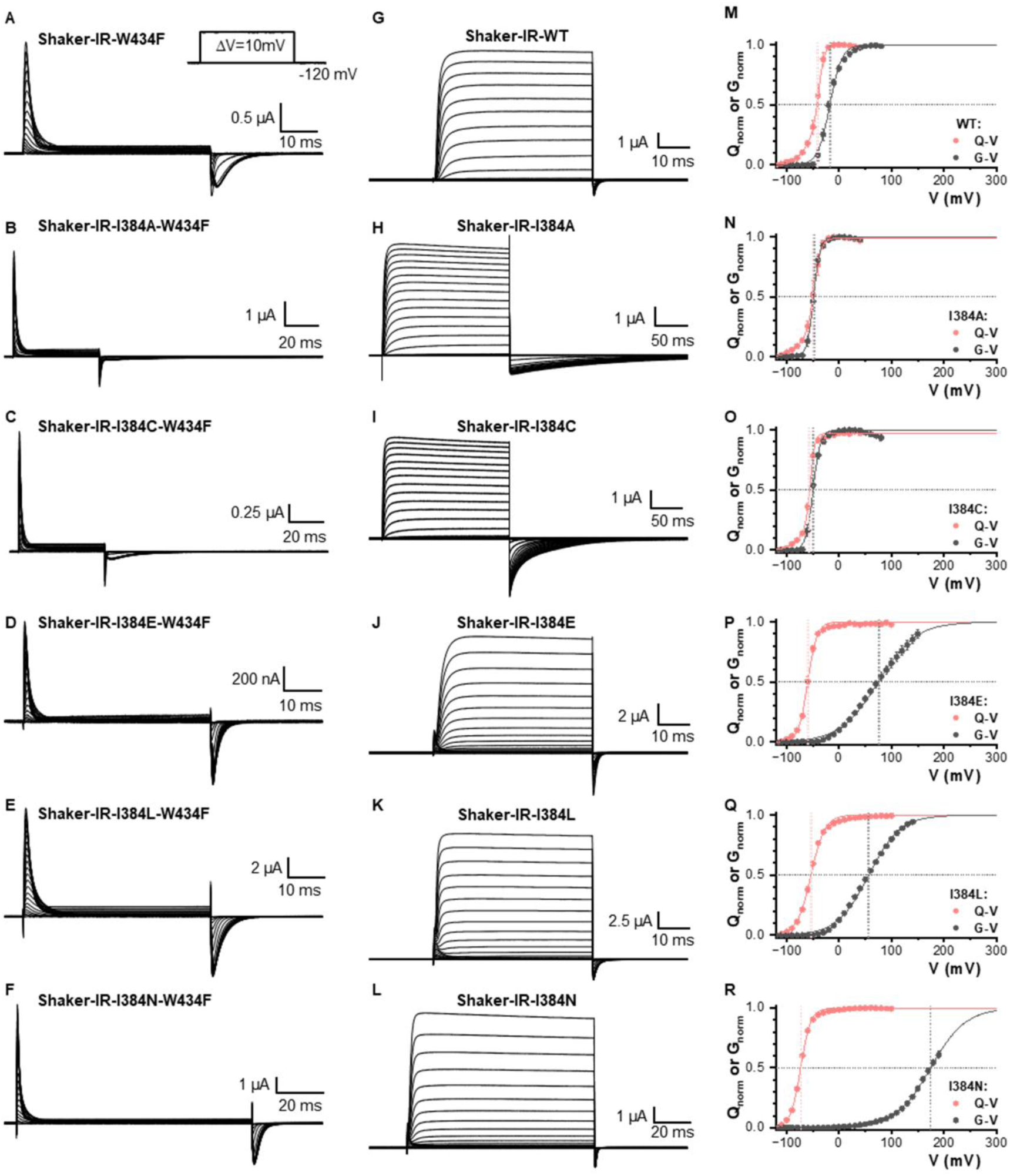
Gating current, ionic current and QV-GV curve comparison for all the I384 mutants in Shaker-IR potassium channel. Representative gating currents for WT **(A)**, I384A **(B)**, I384C **(C)**, I384E **(D)**, I384L **(E)**, and I384N **(F)**. Representative ionic currents for WT **(G)**, I384A **(H)**, I384C **(I)**, I384E **(J)**, I384L **(K)**, and I384N **(L)**. Note that the gating current experiments were conducted with W434F background. The comparison of the QV/GV WT **(M)** curve indicates that I384A **(N)** and I384C **(O)** strengthen the coupling while I384E **(P)**, I384L **(Q)** and I384N **(R)** weaken the coupling.

**Supplementary Figure 2.**
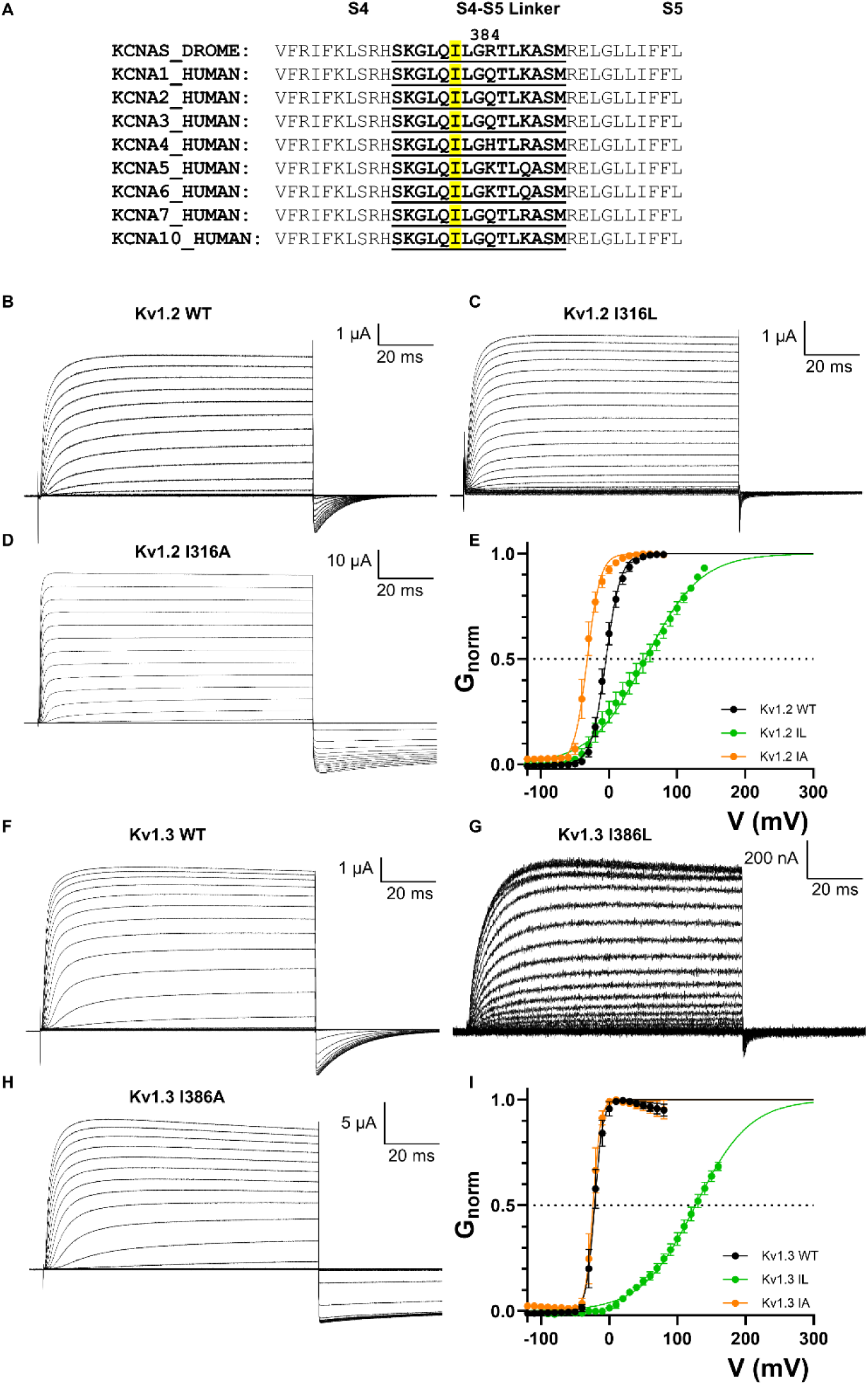
Conserved Isoleucine that controls EMC in Kv1 families. **A)** Sequence alignment of the Shaker potassium channel with human Kv1 channels. I384 is conserved across all the channels. **B)-D)** Results from hKv1.2 mutating the equivalent isoleucine. **F)-I)** Results from hKv1.3 mutating the equivalent isoleucine. Similar behaviors were seen in both channels.

**Supplementary Figure 3.**
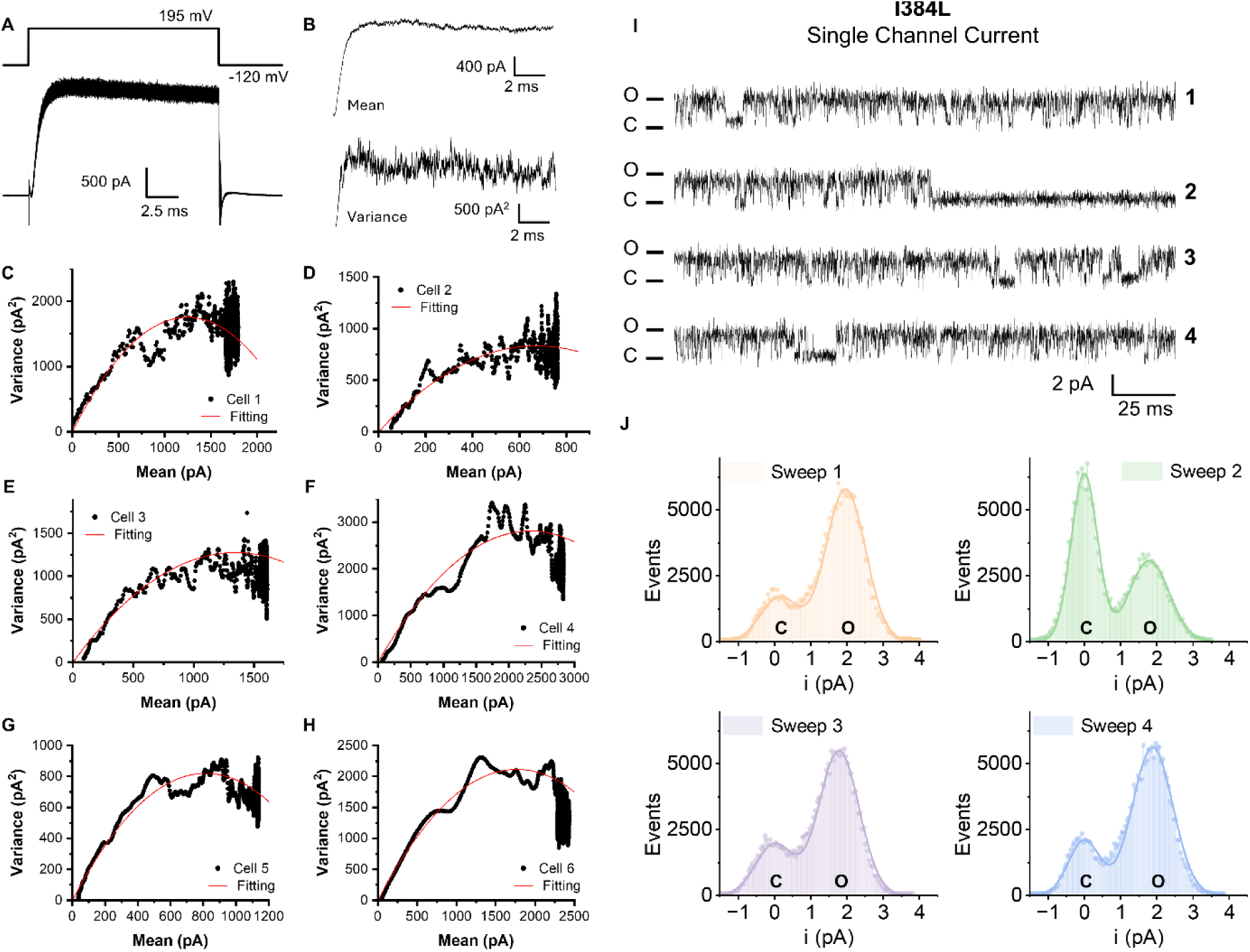
Noise analysis and Single channel recording of I384L. Six independent noise analysis experiments were shown with their fitting parameters **(A-H)**. Voltage protocol is inset in **A**. Representative 150 current traces overlapped are shown in **A)** and in **B)** is depicted the Mean and variance from those recordings. **I)** Four representative current traces for single channel records at 195mV, and increased flickering behavior was observed. **J)** Histogram of the single channel recording traces shown in **I)**.

**Supplementary Figure 4.**
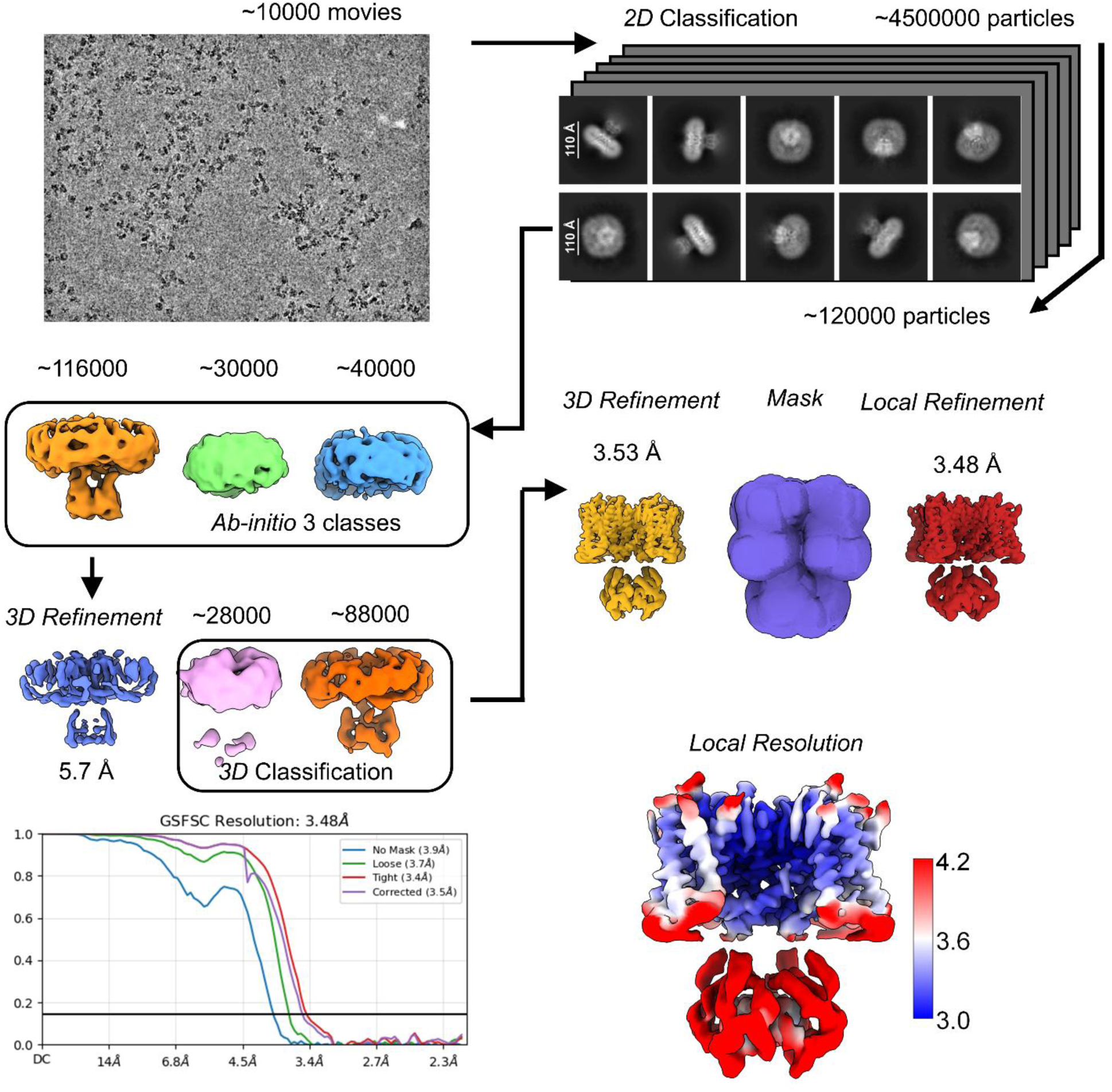
Pipeline of the structural determination for Shaker-IR-I384R. Resolution curve and local resolution map.

**Supplementary Figure 5.**
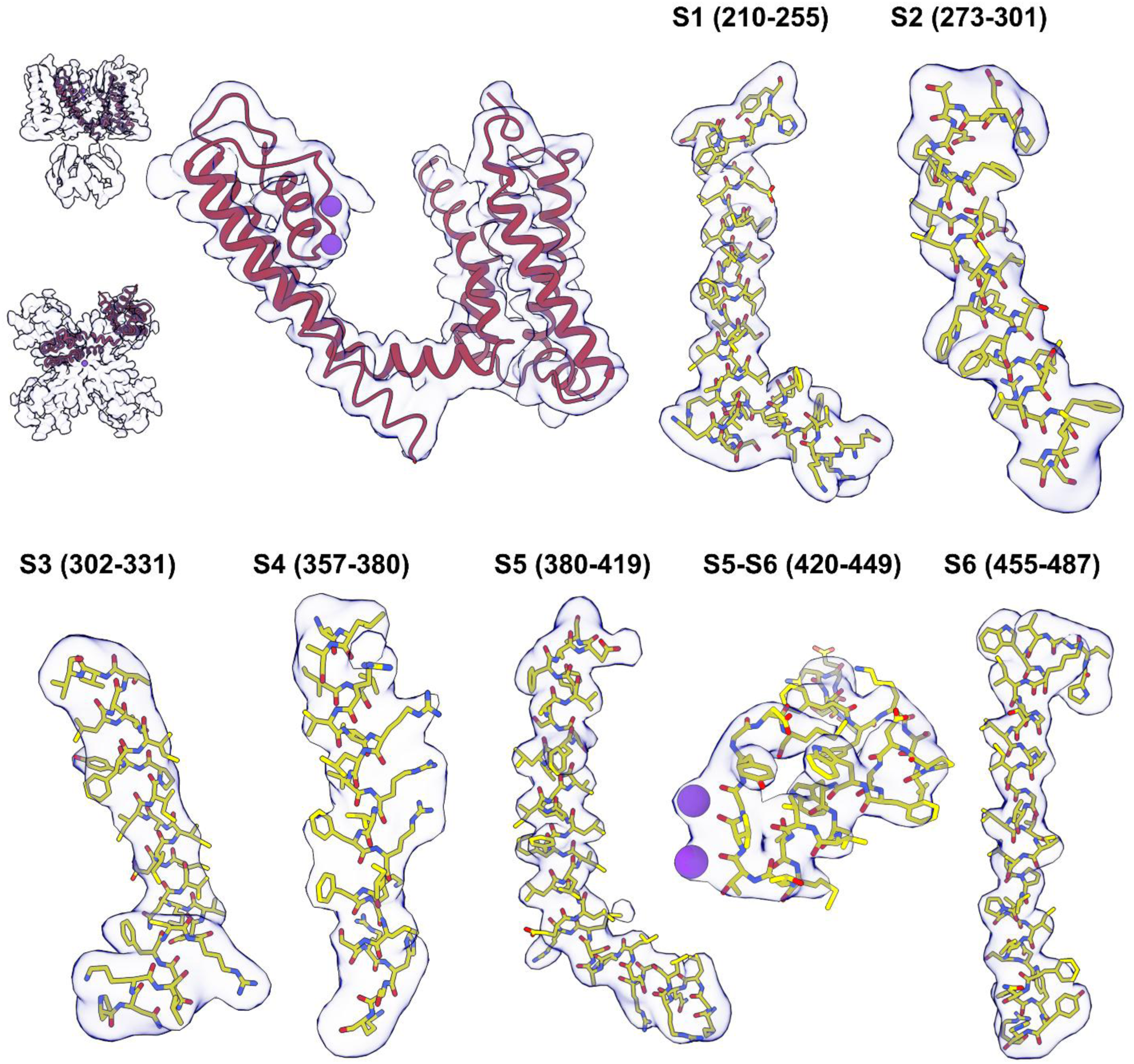
Comparison of atomic model and the density.

**Supplementary Figure 6.**
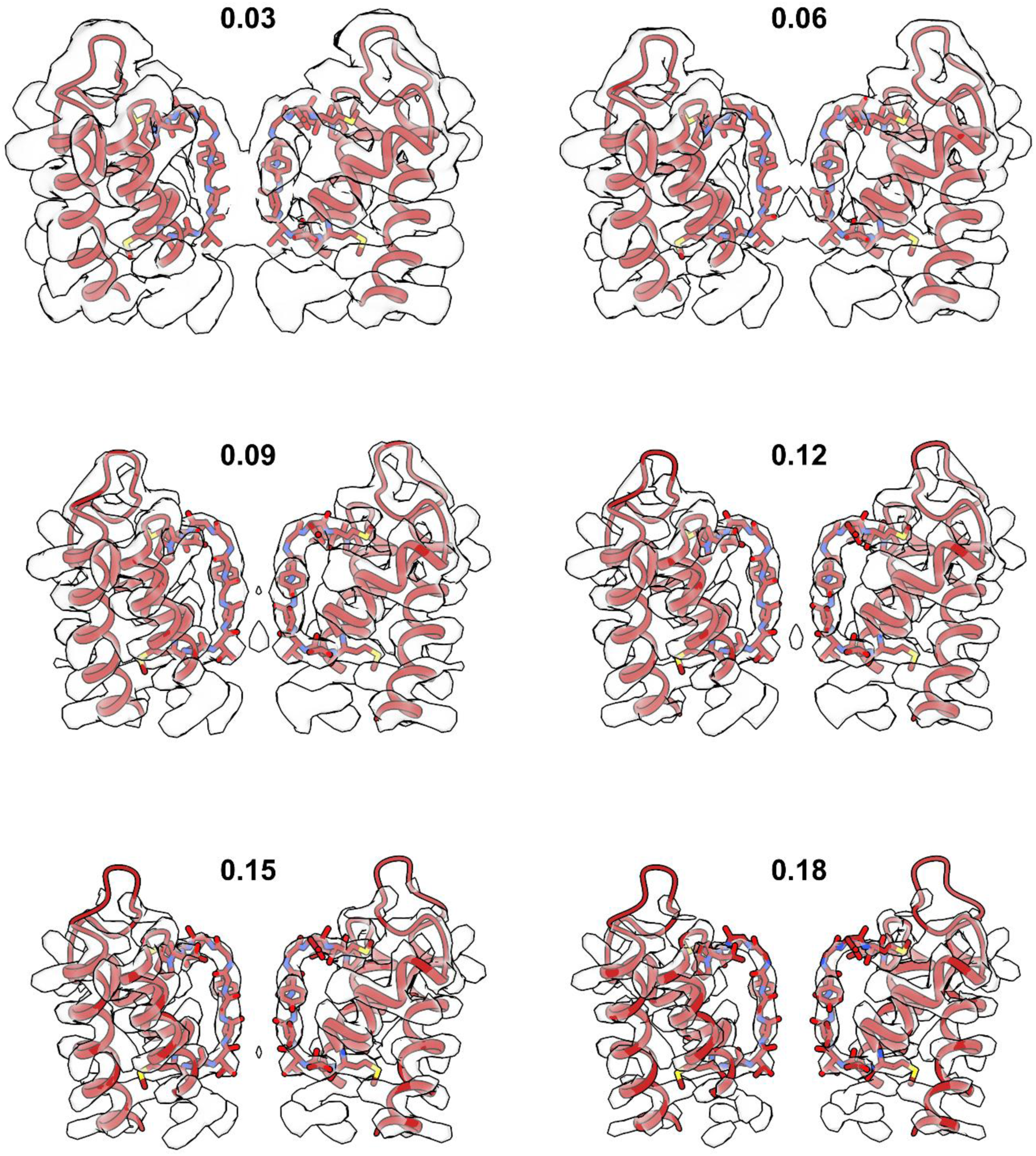
Atomic model overlaid with the density map plotted with variable iso-surface value.

**Supplementary Figure 7.**
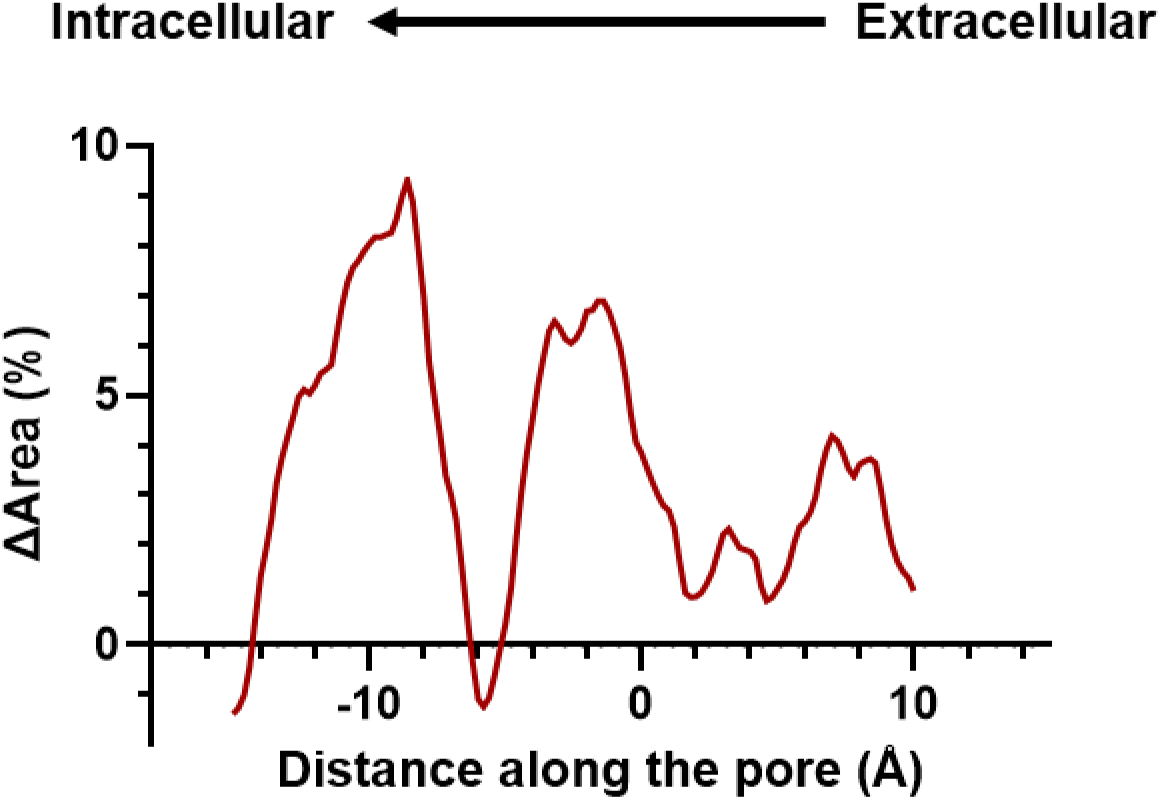
Percent area increase the open state compared to the closed state along the membrane. Positive value indicates expansion in open state.

**Supplementary Figure 8.**
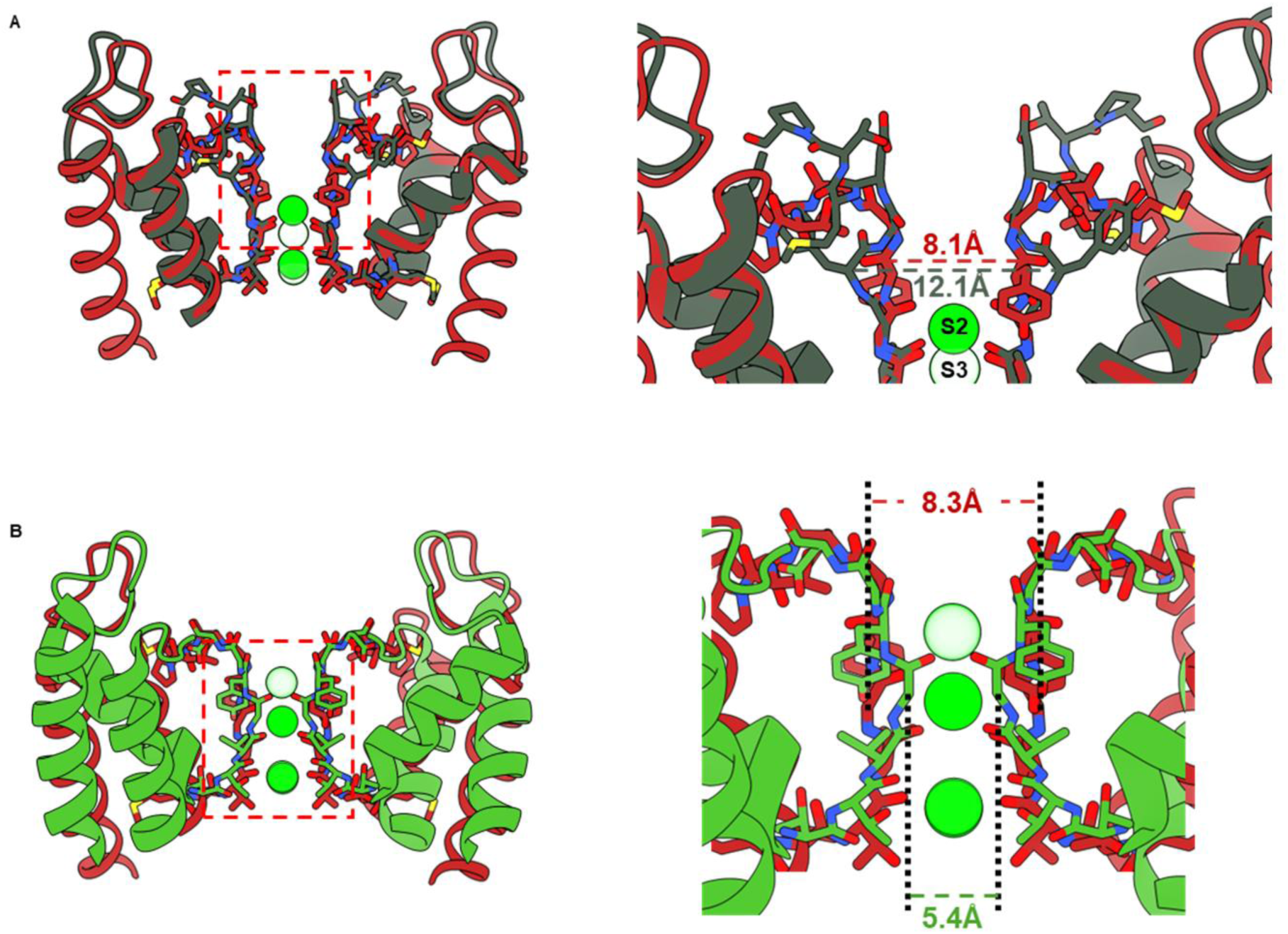
Comparison of the noncanonical filter with dilated filter, **A), B)** (PDB:8ETO) and pinched filter, **C), D)** (PDB:3F5W). The noncanonical filter is different from both conformations.

**Supplementary Figure 9.**
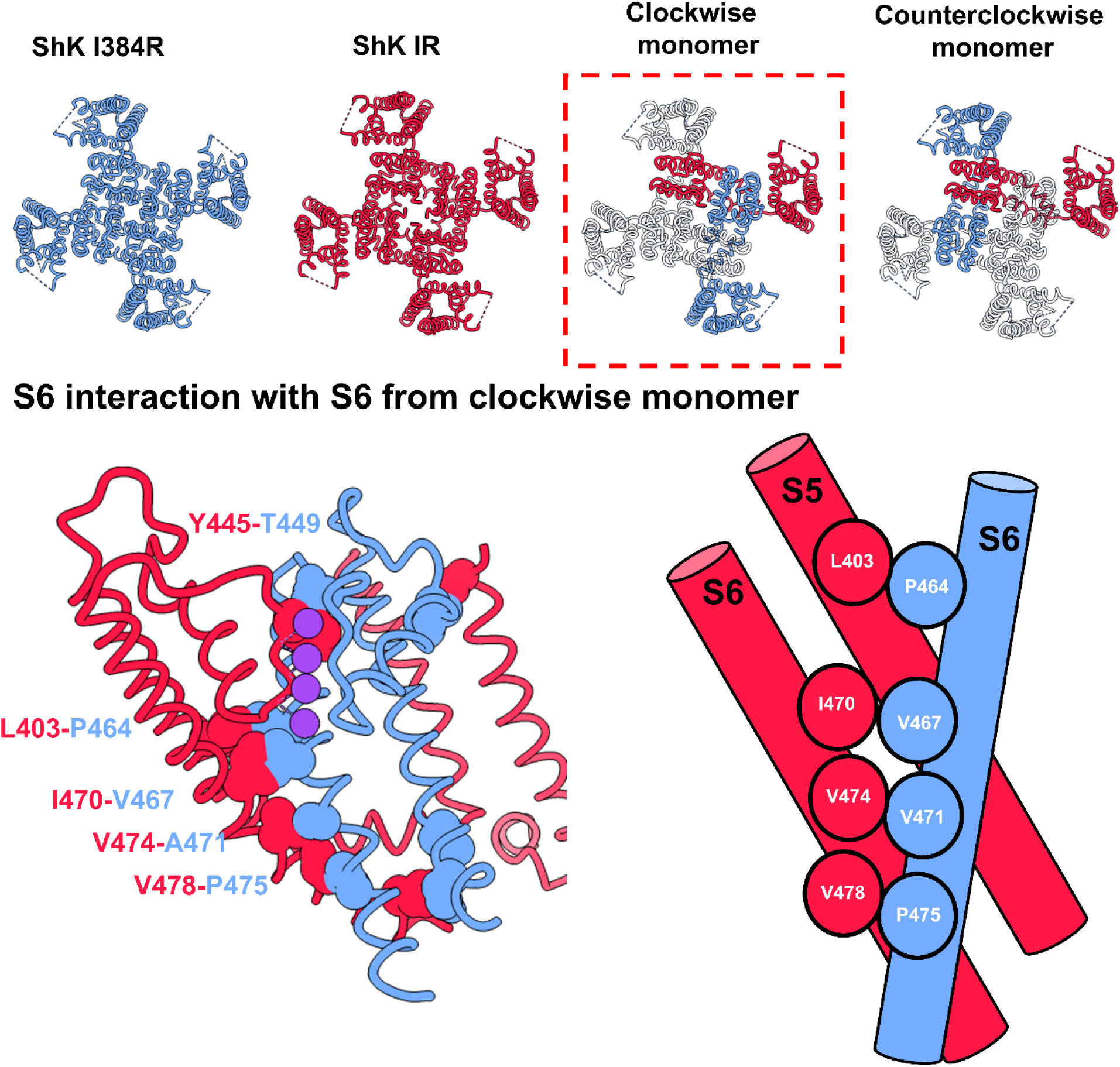
Clash analysis by assembling open state and closed state structure into the same homotetramer. The lower panel shows the pair clashes happened between the clockwise monomer of open and closed state. Those pairs of residues likely are involved in the activation of the pore.

**Supplementary Figure 10.**
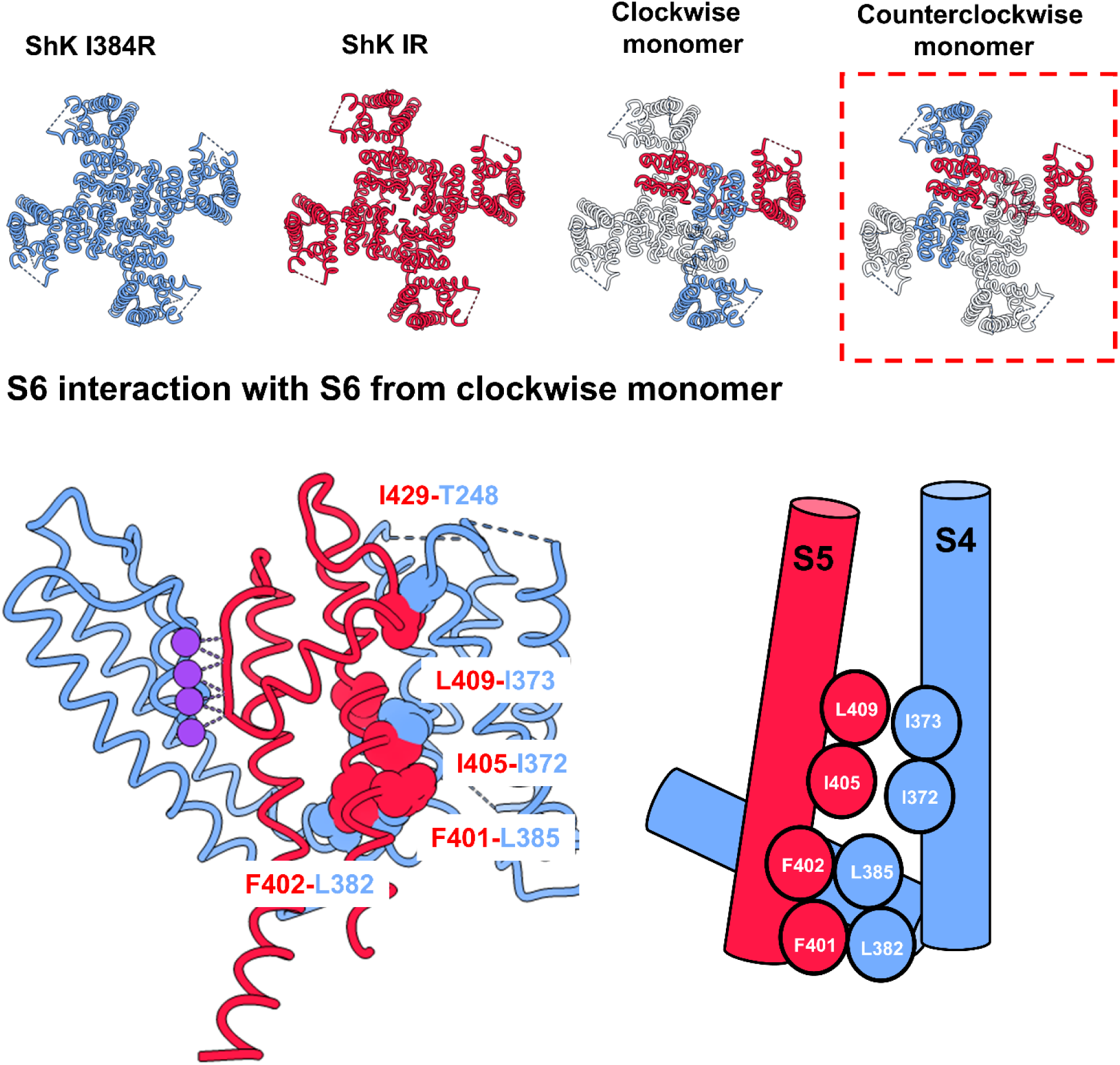
Clash analysis by assembling open state and closed state structure into the same homotetramer. The lower panel shows the pair clashes happened between the counterclockwise monomer of open and closed state. Those pairs of residues likely are involved in the electromechanical coupling or the noncanonical coupling.

## Supplementary Tables

**Table 1:** Fitting parameters for GV curves.

| | $V_{1/2}$ | | | $z$ | | |
| --- | --- | --- | --- | --- | --- | --- |
| ***WT*** | -17.70 | ± | 0.73 | 2.22 | ± | 0.13 |
| ***I384A*** | -48.70 | ± | 0.26 | 4.01 | ± | 0.15 |
| ***I384C*** | -49.80 | ± | 0.62 | 3.64 | ± | 0.29 |
| ***I384E*** | 74.45 | ± | 0.91 | 0.70 | ± | 0.02 |
| ***I384L*** | 56.12 | ± | 0.70 | 0.84 | ± | 0.02 |
| ***I384N*** | 174.33 | ± | 0.25 | 0.76 | ± | 0.01 |

**Table 2:**
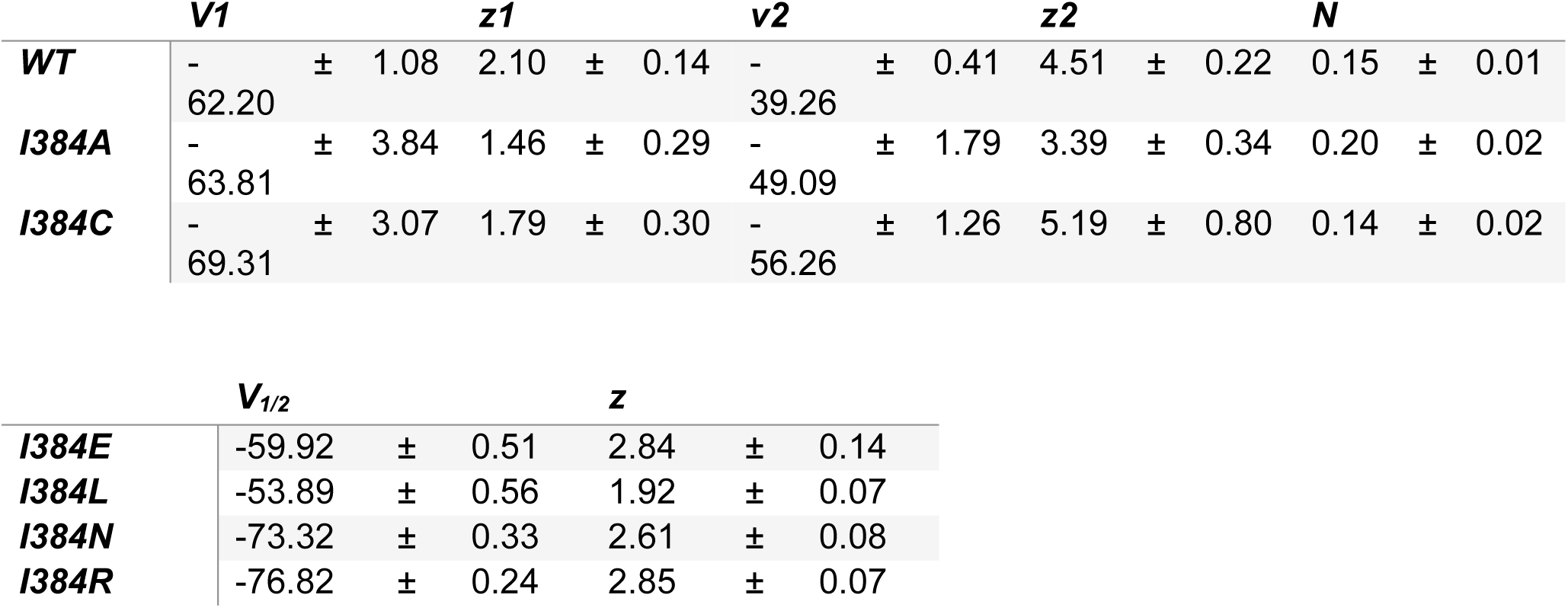
Fitting parameters for QV curves.

**Table 3:**
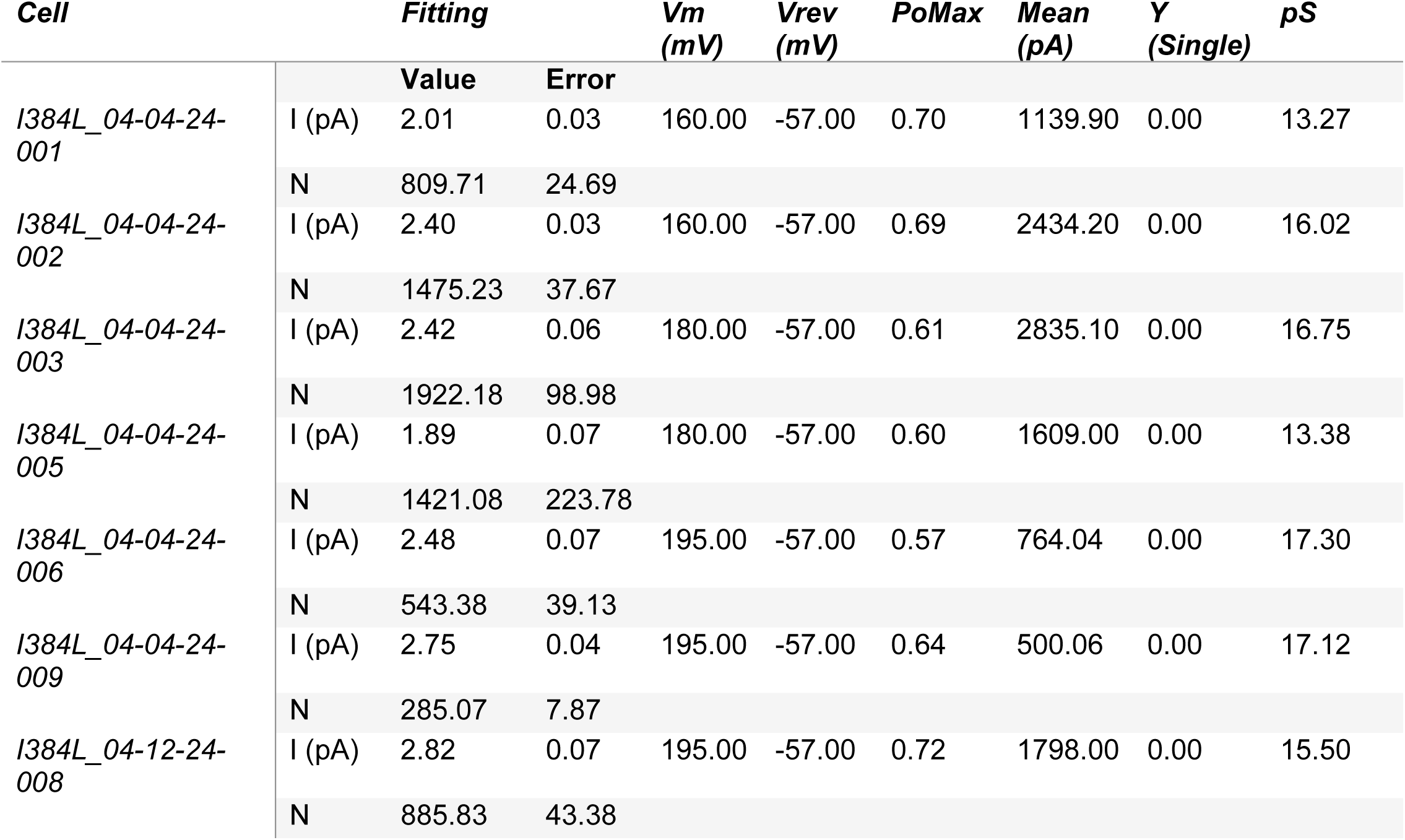
Fitting parameters for noise analysis.

**Table 4:** Fitting parameters for single channel analysis.

|  | *Sweep 1* |  | *Sweep 2* |  | *Sweep 3* |  | *Sweep 4* |  |
| --- | --- | --- | --- | --- | --- | --- | --- | --- |
|  | **Value** | **error** | **Value** | **error** | **Value** | **error** | **Value** | **error** |
| **$\mu 1$** | 3.93 | 0.01 | 3.75 | 0.01 | 3.94 | 0.01 | 3.99 | 0.01 |
| **$\sigma 1$** | 0.77 | 0.01 | 0.78 | 0.01 | 0.78 | 0.01 | 0.79 | 0.01 |
| **$A1$** | 22399.67 | 243.33 | 11969.20 | 182.66 | 21546.90 | 235.23 | 22217.59 | 255.66 |
| **$\mu 2$** | 2.11 | 0.02 | 1.97 | 0.00 | 2.16 | 0.02 | 2.11 | 0.02 |
| **$\sigma 2$** | 0.73 | 0.03 | 0.51 | 0.01 | 0.73 | 0.03 | 0.62 | 0.03 |
| **$A2$** | 6187.80 | 238.43 | 16331.76 | 150.59 | 7126.41 | 229.39 | 6420.97 | 228.52 |

## References

Afonine, P. V., Poon, B. K., Read, R. J., Sobolev, O. V., Terwilliger, T. C., Urzhumtsev, A., & Adams, P. D. (2018). Real-space refinement in PHENIX for cryo-EM and crystallography. Acta Crystallographica. Section D, Structural Biology, 74(Pt 6), 531–544. 10.1107/S2059798318006551

Aggarwal, S. K., & MacKinnon, R. (1996). Contribution of the S4 Segment to Gating Charge in the Shaker K+ Channel. Neuron, 16(6), 1169–1177. 10.1016/S0896-6273(00)80143-9

Alvarez, O., Gonzalez, C., & Latorre, R. (2002). Counting channels: A tutorial guide on ion channel fluctuation analysis. American Journal of Physiology - Advances in Physiology Education, 26(1–4), 327–341. 10.1152/advan.00006.2002

Armstrong, C. M. (1971). Interaction of tetraethylammonium ion derivatives with the potassium channels of giant axons. The Journal of General Physiology, 58(4), 413–437. 10.1085/JGP.58.4.413

Armstrong, C. M., & Bezanilla, F. (1977). Inactivation of the sodium gating current. The Journal of General Physiology, 4(7), 865–876. 10.1016/0306-4522(79)90171-4

Bassetto, C. A. Z., Costa, F., Guardiani, C., Bezanilla, F., & Giacomello, A. (2023). Noncanonical electromechanical coupling paths in cardiac hERG potassium channel. Nature Communications, 14(1), 1110. 10.1038/s41467-023-36730-7

Batulan, Z., Haddad, G. A., & Blunck, R. (2010). An Intersubunit Interaction between S4-S5 Linker and S6 Is Responsible for the Slow Off-gating Component in Shaker K+ Channels. The Journal of Biological Chemistry, 285(18), 14005. 10.1074/JBC.M109.097717

Bezanilla, F. (2018). Gating currents. Journal of General Physiology, 150(7), 911–932. 10.1085/jgp.201812090

Bezanilla, F., Taylor, R. E., & Fernández, J. M. (1982). Distribution and kinetics of membrane dielectric polarization: 1. long-term inactivation of gating currents. Journal of General Physiology, 79(1), 21–40. 10.1085/jgp.79.1.21

Cha, A., & Bezanilla, F. (1997). Characterizing Voltage-Dependent Conformational Changes in the ShakerK+ Channel with Fluorescence. Neuron, 19(5), 1127–1140. 10.1016/S0896-6273(00)80403-1

Chatterjee, A., Guo, J., Lee, H. S., & Schultz, P. G. (2013). A genetically encoded fluorescent probe in mammalian cells. Journal of the American Chemical Society, 135(34), 12540–12543. 10.1021/ja4059553

Chowdhury, S., Haehnel, B. M., & Chanda, B. (2014). Interfacial gating triad is crucial for electromechanical transduction in voltage-activated potassium channels. The Journal of General Physiology, 144(5), 457–467. 10.1085/JGP.201411185

Colquhoun, D., & Sigworth, F. J. (1995). Fitting and Statistical Analysis of Single-Channel 10.1007/978-1-4419-1229-9_19

Croll, T. I. (2018). ISOLDE: a physically realistic environment for model building into low-resolution electron-density maps. Acta Crystallographica. Section D, Structural Biology, 74(Pt 6), 519–530. 10.1107/S2059798318002425

Cuello, L. G., Cortes, D. M., & Perozo, E. (2017). The gating cycle of a K+ channel at atomic resolution. ELife, 6, 1–17. 10.7554/eLife.28032

Cuello, L. G., Jogini, V., Cortes, D. M., & Perozo, E. (2010). Structural mechanism of C-type inactivation in K(+) channels. Nature, 466(7303), 203–208. 10.1038/NATURE09153

Dalal, V., Arcario, M. J., Petroff, J. T., Tan, B. K., Dietzen, N. M., Rau, M. J., Fitzpatrick, J. A. J., Brannigan, G., & Cheng, W. W. L. (2024). Lipid nanodisc scaffold and size alter the structure of a pentameric ligand-gated ion channel. Nature Communications 2024 15:1, 15(1), 1–10. 10.1038/s41467-023-44366-w

Del Camino, D., Holmgren, M., Liu, Y., & Yellen, G. (2000). Blocker protection in the pore of a voltage-gated K+ channel and its structural implications. Nature 2000 403:6767, 403(6767), 321–325. 10.1038/35002099

Del Camino, D., & Yellen, G. (2001). Tight steric closure at the intracellular activation gate of a voltage-gated K+ channel. Neuron, 32(4), 649–656. 10.1016/S0896-6273(01)00487-1

Ding, S., & Horn, R. (2003). Effect of S6 Tail Mutations on Charge Movement in Shaker Potassium Channels. Biophysical Journal, 84(1), 295. 10.1016/S0006-3495(03)74850-4

Doyle, D. A., Cabral, J. M., Pfuetzner, R. A., Kuo, A., Gulbis, J. M., Cohen, S. L., Chait, B. T., & MacKinnon, R. (1998). The structure of the potassium channel: molecular basis of K+ conduction and selectivity. Science (New York, N.Y.), 280(5360), 69–77. 10.1126/SCIENCE.280.5360.69

Durner, A., Durner, E., & Nicke, A. (2023). Improved ANAP incorporation and VCF analysis reveal details of P2X7 current facilitation and a limited conformational interplay between ATP binding and the intracellular ballast domain. ELife, 12, 1–32. 10.7554/elife.82479

Emsley, P., Lohkamp, B., Scott, W. G., & Cowtan, K. (2010). Features and development of Coot. Acta Crystallographica. Section D, Biological Crystallography, 66(Pt 4), 486–501. 10.1107/S0907444910007493

Flynn, G. E., & Zagotta, W. N. (2018). Insights into the molecular mechanism for hyperpolarization-dependent activation of HCN channels. Proceedings of the National Academy of Sciences of the United States of America, 115(34), E8086– E8095. 10.1073/pnas.1805596115

Fowler, P. W., & Sansom, M. S. P. (2013). The pore of voltage-gated potassium ion channels is strained when closed. Nature Communications 2013 4:1, 4(1), 1–8.

Goddard, T. D., Huang, C. C., Meng, E. C., Pettersen, E. F., Couch, G. S., Morris, J. H., & Ferrin, T. E. (2018). UCSF ChimeraX: Meeting modern challenges in visualization and analysis. Protein Science : A Publication of the Protein Society, 27(1), 14–25. 10.1002/PRO.3235

Hackos, D. H., Chang, T. H., & Swartz, K. J. (2002). Scanning the Intracellular S6 Activation Gate in the Shaker K+ Channel. The Journal of General Physiology, 119(6), 521. 10.1085/JGP.20028569

Haddad, G. A., & Blunck, R. (2011). Mode shift of the voltage sensors in Shaker K+ channels is caused by energetic coupling to the pore domain. The Journal of General Physiology, 137(5), 455–472. 10.1085/JGP.201010573

Hirschberg, B., Rovner, A., Lieberman, M., & Patlak, J. (1995). Transfer of twelve charges is needed to open skeletal muscle Na+ channels. Journal of General Physiology, 106(6), 1053–1068. 10.1085/jgp.106.6.1053

Holmgren, M., Shin, K. S., & Yellen, G. (1998). The activation gate of a voltage-gated K+ channel can be trapped in the open state by an intersubunit metal bridge. Neuron, 21(3), 617–621. 10.1016/S0896-6273(00)80571-1

Holmgren, M., Smith, P. L., & Yellen, G. (1997). Trapping of organic blockers by closing of voltage-dependent K+ channels: Evidence for a trap door mechanism of activation gating. Journal of General Physiology, 109(5), 527–535. 10.1085/jgp.109.5.527

Hoshi, T., Zagotta, W. N., & Aldrich, R. W. (1990). Biophysical and Molecular Mechanisms of Shaker Potassium Channel Inactivation. Science, 250(4980), 533–538. 10.1126/science.2122519

Hyun, S. L., Guo, J., Lemke, E. A., Dimla, R. D., & Schultz, P. G. (2009). Genetic incorporation of a small, environmentally sensitive, fluorescent probe into proteins in Saccharomyces cerevisiae. Journal of the American Chemical Society, 131(36), 12921–12923. 10.1021/ja904896s

Isacoff, E. Y., Jan, Y. N., & Jan, L. Y. (1991). Putative receptor for the cytoplasmic inactivation gate in the Shaker K+ channel. Nature 1991 353:6339, 353(6339), 86–90. 10.1038/353086a0

Islas, L. D., & Sigworth, F. J. (1999). Voltage Sensitivity and Gating Charge in Shaker and Shab Family Potassium Channels. Journal of General Physiology, 114(5), 723– 742. 10.1085/JGP.114.5.723

Jan, L. Y., & Jan, Y. N. (2012). Voltage-gated potassium channels and the diversity of electrical signalling. The Journal of Physiology, 590(11), 2591–2599. 10.1113/JPHYSIOL.2011.224212

Jo, S., Kim, T., Iyer, V. G., & Im, W. (2008). CHARMM-GUI: A web-based graphical user interface for CHARMM. Journal of Computational Chemistry, 29(11), 1859–1865. 10.1002/JCC.20945

Kalstrup, T., & Blunck, R. (2013). Dynamics of internal pore opening in KV channels probed by a fluorescent unnatural amino acid. Proceedings of the National Academy of Sciences of the United States of America, 110(20), 8272–8277. 10.1073/pnas.1220398110

Kalstrup, T., & Blunck, R. (2017). Voltage-clamp fluorometry in Xenopus oocytes using fluorescent unnatural amino acids. Journal of Visualized Experiments, 2017(123), 1–9. 10.3791/55598

Lacroix, J. J., & Bezanilla, F. (2011). Control of a final gating charge transition by a hydrophobic residue in the S2 segment of a K+ channel voltage sensor. Proceedings of the National Academy of Sciences of the United States of America, 108(16), 6444– 6449. 10.1073/pnas.1103397108

Lacroix, J. J., Hyde, H. C., Campos, F. V, & Bezanilla, F. (2014). Moving gating charges through the gating pore in a Kv channel voltage sensor. Proceedings of the National Academy of Sciences, 111(19), E1950–9. 10.1073/pnas.1406161111

Lacroix, J. J., Labro, A. J., & Bezanilla, F. (2011). Properties of Deactivation Gating Currents in Shaker Channels. Biophysical Journal, 100(5), L28–L30. 10.1016/J.BPJ.2011.01.043

Laitko, U., & Morris, C. E. (2004). Membrane Tension Accelerates Rate-limiting Voltage-dependent Activation and Slow Inactivation Steps in a Shaker Channel. The Journal of General Physiology, 123(2), 135. 10.1085/JGP.200308965

Latorre, R., Morera, F. J., & Zaelzer, C. (2010). SYMPOSIUM REVIEW: Allosteric interactions and the modular nature of the voltage- and Ca 2+ -activated (BK) channel. The Journal of Physiology, 588(17), 3141–3148. 10.1113/jphysiol.2010.191999

Lee, B. R., White, K. I., Socolich, M., Klureza, M. A., Henning, R., Srajer, V., Ranganathan, R., & Hekstra, D. R. (2025). Direct visualization of electric-field-stimulated ion conduction in a potassium channel. Cell, 188(1), 77–88.e15. 10.1016/J.CELL.2024.12.006/ATTACHMENT/2B6E36D7-7F0A-4B35-9D53-63F52AAD8909/MMC5.PDF

Liu, Y., Bassetto, C. A. Z., Pinto, B. I., & Bezanilla, F. (2023). A mechanistic reinterpretation of fast inactivation in voltage-gated Na+ channels. Nature Communications 2023 14:1, 14(1), 1–13. 10.1038/s41467-023-40514-4

Liu, Y., Holmgren, M., Jurman, M. E., & Yellen, G. (1997). Gated access to the pore of a voltage-dependent K+ channel. Neuron, 19(1), 175–184. 10.1016/S0896-6273(00)80357-8

Long, S. B., Tao, X., Campbell, E. B., & MacKinnon, R. (2007). Atomic structure of a voltage-dependent K+ channel in a lipid membrane-like environment. Nature 2007 450:7168, 450(7168), 376–382. 10.1038/nature06265

Lörinczi, É., Gómez-Posada, J. C., de la Peña, P., Tomczak, A. P., Fernández-Trillo, J., Leipscher, U., Stühmer, W., Barros, F., & Pardo, L. A. (2015). Voltage-dependent gating of KCNH potassium channels lacking a covalent link between voltage-sensing and pore domains. Nature Communications, 6(1), 6672. 10.1038/ncomms7672

Mannuzzu, L. M., Moronne, M. M., & Isacoff, E. Y. (1996). Direct physical measure of conformational rearrangement underlying potassium channel gating. Science (New York, N.Y.), 271(5246), 213–216. 10.1126/SCIENCE.271.5246.213

McCormack, K., Tanouye, M. A., Iverson, L. E., Lin, J. W., Ramaswami, M., McCormack, T., Campanelli, J. T., Mathew, M. K., & Rudy, B. (1991). A role for hydrophobic residues in the voltage-dependent gating of Shaker K+ channels. Proceedings of the National Academy of Sciences, 88(7), 2931–2935. 10.1073/PNAS.88.7.2931

Morris, C. E. (2011). Voltage-gated channel mechanosensitivity: Fact or friction? Frontiers in Physiology, MAY(May), 1–10. 10.3389/fphys.2011.00025

Morris, C. E., & Juranka, P. F. (2007). Nav Channel Mechanosensitivity: Activation and Inactivation Accelerate Reversibly with Stretch. Biophysical Journal, 93(3), 822. 10.1529/BIOPHYSJ.106.101246

Olcese, R., Latorre, R., Toro, L., Bezanilla, F., & Stefani, E. (1997). Correlation between charge movement and ionic current during slow inactivation in Shaker K+ channels. The Journal of General Physiology, 110(5), 579–589. 10.1085/JGP.110.5.579

Olcese, R., Sigg, D., Latorre, R., Bezanilla, F., & Stefani, E. (2001). A Conducting State with Properties of a Slow Inactivated State in a Shaker K+ Channel Mutant. The Journal of General Physiology, 117(2), 149. 10.1085/JGP.117.2.149

Panyi, G., & Deutsch, C. (2006). Cross Talk between Activation and Slow Inactivation Gates of Shaker Potassium Channels. 128(5), 547–559. 10.1085/jgp.200609644

Panyi, G., & Deutsch, C. (2007). Probing the Cavity of the Slow Inactivated Conformation of Shaker Potassium Channels. Journal of General Physiology, 129(5), 403–418. 10.1085/JGP.200709758

Perozo, E., MacKinnon, R., Bezanilla, F., & Stefani, E. (1993). Gating currents from a nonconducting mutant reveal open-closed conformations in Shaker K+channels. Neuron, 11(2), 353–358. 10.1016/0896-6273(93)90190-3

Peters, C. J., Fedida, D., & Accili, E. A. (2013). Allosteric coupling of the inner activation gate to the outer pore of a potassium channel. Scientific Reports 2013 3:1, 3(1), 1–8. 10.1038/srep03025

Pettersen, E. F., Goddard, T. D., Huang, C. C., Meng, E. C., Couch, G. S., Croll, T. I., Morris, J. H., & Ferrin, T. E. (2021). UCSF ChimeraX: Structure visualization for researchers, educators, and developers. Protein Science, 30(1), 70–82. 10.1002/pro.3943

Punjani, A., Rubinstein, J. L., Fleet, D. J., & Brubaker, M. A. (2017). cryoSPARC: algorithms for rapid unsupervised cryo-EM structure determination. Nature Methods, 14(3), 290–296. 10.1038/NMETH.4169

Punjani, A., Zhang, H., & Fleet, D. J. (2020). Non-uniform refinement: adaptive regularization improves single-particle cryo-EM reconstruction. Nature Methods, 17(12), 1214–1221. 10.1038/S41592-020-00990-8

Schmidt, D., Del Maŕmol, J., & MacKinnon, R. (2012). Mechanistic basis for low threshold mechanosensitivity in voltage-dependent K+ channels. Proceedings of the National Academy of Sciences of the United States of America, 109(26), 10352–10357. 10.1073/PNAS.1204700109/SUPPL_FILE/SM01.MOV

Seoh, S. A., Sigg, D., Papazian, D. M., & Bezanilla, F. (1996). Voltage-sensing residues in the S2 and S4 segments of the Shaker K+ channel. Neuron, 16(6), 1159–1167. 10.1016/S0896-6273(00)80142-7

Shirokov, R., Levis, R., Shirokova, N., & Ríos, E. (1992). Two classes of gating current from L-type Ca channels in guinea pig ventricular myocytes. Journal of General Physiology, 99(6), 863–895. 10.1085/JGP.99.6.863

Sigg, D., & Bezanilla, F. (1997). Total charge movement per channel: The relation between gating charge displacement and the voltage sensitivity of activation. Journal of General Physiology, 109(1), 27–39. 10.1085/jgp.109.1.27

Sigworth, F. J. (1980). The variance of sodium current fluctuations at the node of Ranvier. The Journal of Physiology, 307(1), 97. 10.1113/JPHYSIOL1980.SP013426

Slesinger, P. A., Jan, Y. N., & Jan, L. Y. (1993). The S4–S5 loop contributes to the ion-selective pore of potassium channels. Neuron, 11(4), 739–749. 10.1016/0896-6273(93)90083-4

Smart, O. S., Neduvelil, J. G., Wang, X., Wallace, B. A., & Sansom, M. S. P. (1996). HOLE: A program for the analysis of the pore dimensions of ion channel structural models. Journal of Molecular Graphics, 14(6), 354–360. 10.1016/S0263-7855(97)00009-X

Stefani, E., & Bezanilla, F. (1998). Cut-open oocyte voltage-clamp technique. Methods in Enzymology, 293(1987), 300–318. 10.1016/S0076-6879(98)93020-8

Stix, R., Tan, X. F., Bae, C., Fernández-Mariño, A. I., Swartz, K. J., & Faraldo-Gómez, J. D. (2023). Eukaryotic Kv channel Shaker inactivates through selectivity filter dilation rather than collapse. Science Advances, 9(49). 10.1126/SCIADV.ADJ5539/SUPPL_FILE/SCIADV.ADJ5539_MOVIES_S1_TO_S5.ZIP

Szanto, T. G., Zakany, F., Papp, F., Varga, Z., Deutsch, C. J., & Panyi, G. (2020). The activation gate controls steady-state inactivation and recovery from inactivation in Shaker. Journal of General Physiology, 152(8). 10.1085/jgp.202012591

Tabarean, I. V., Juranka, P., & Morris, C. E. (1999). Membrane stretch affects gating modes of a skeletal muscle sodium channel. Biophysical Journal, 77(2), 758–774. 10.1016/S0006-3495(99)76930-4

Tan, X. F., Bae, C., Stix, R., Fernández-Mariño, A. I., Huffer, K., Chang, T. H., Jiang, J., Faraldo-Gómez, J. D., & Swartz, K. J. (2022). Structure of the Shaker Kv channel and mechanism of slow C-type inactivation. Science Advances, 8(11). 10.1126/sciadv.abm7814

Tao, X., Lee, A., Limapichat, W., Dougherty, D. a, & MacKinnon, R. (2010). A gating charge transfer center in voltage sensors. Science (New York, N.Y.), 328(5974), 67–73. 10.1126/science.1185954

Tombola, F., Pathak, M. M., & Isacoff, E. Y. (2006). How does voltage open an ion channel? Annual Review of Cell and Developmental Biology, 22, 23–52. 10.1146/annurev.cellbio.21.020404.145837

Treptow, W., Liu, Y., Bassetto, C. A., Pinto, B. I., Alves Nunes, J. A., Uriarte, R. M., Chipot, C. J., Bezanilla, F., & Roux, B. (2024). Isoleucine gate blocks K+ conduction in C-type inactivation. ELife, 13. 10.7554/ELIFE.97696

Villalba-Galea, C. A., Sandtner, W., Starace, D. M., & Bezanilla, F. (2008). S4-based voltage sensors have three major conformations. Proceedings of the National Academy of Sciences of the United States of America, 105(46), 17600–17607. 10.1073/PNAS.0807387105/SUPPL_FILE/0807387105SI.PDF

Webster, S. M., Del Camino, D., Dekker, J. P., & Yellen, G. (2004). Intracellular gate opening in Shaker K+ channels defined by high-affinity metal bridges. Nature, 428(6985), 864–868. 10.1038/nature02468

Williams, C. J., Headd, J. J., Moriarty, N. W., Prisant, M. G., Videau, L. L., Deis, L. N., Verma, V., Keedy, D. A., Hintze, B. J., Chen, V. B., Jain, S., Lewis, S. M., Arendall, W. B., Snoeyink, J., Adams, P. D., Lovell, S. C., Richardson, J. S., & Richardson, D. C. (2018). MolProbity: More and better reference data for improved all-atom structure validation. Protein Science, 27(1), 293–315. 10.1002/PRO.3330

Wisedchaisri, G., Tonggu, L., McCord, E., Gamal El-Din, T. M., Wang, L., Zheng, N., & Catterall, W. A. (2019). Resting-State Structure and Gating Mechanism of a Voltage-Gated Sodium Channel. Cell, 178(4), 993–1003.e12. 10.1016/j.cell.2019.06.031

Yang, Y., Yan, Y., & Sigworth, F. J. (1997). How Does the W434F Mutation Block Current in Shaker Potassium Channels? The Journal of General Physiology, 109(6), 779. 10.1085/JGP.109.6.779

Ye, W., Zhao, H., Dai, Y., Wang, Y., Lo, Y., Jan, L. Y., & Lee, C.-H. (2022). Activation and closed-state inactivation mechanisms of the human voltage-gated KV4 channel complexes. Molecular Cell, 1–16. 10.1016/j.molcel.2022.04.032

Yifrach, O., & Mackinnon, R. (2002). Energetics of Pore Opening in a Voltage-Gated K ؉ Channel. 111, 231–239.

Zagotta, W. N., Hoshi, T., & Aldrich, R. W. (1994). Shaker potassium channel gating. III: Evaluation of kinetic models for activation. Journal of General Physiology, 103(2), 321–362. 10.1085/JGP.103.2.321

